# Global biogeography of N_2_-fixing microbes: *nifH* amplicon database and analytics workflow

**DOI:** 10.1101/2024.05.04.592440

**Authors:** Michael Morando, Jonathan Magasin, Shunyan Cheung, Matthew M. Mills, Jonathan P. Zehr, Kendra A. Turk-Kubo

**Affiliations:** Ocean Sciences Department, University of California, Santa Cruz, Santa Cruz, 95064, United States; Institute of Marine Biology and Center of Excellence for the Oceans, National Taiwan Ocean University, Keelung, Taiwan; Earth System Science, Stanford University, Stanford, 94305, United States

## Abstract

Marine nitrogen (N) fixation is a globally significant biogeochemical process carried out by a specialized group of prokaryotes (diazotrophs), yet our understanding of their ecology is constantly evolving. Although marine dinitrogen (N_2_)-fixation is often ascribed to cyanobacterial diazotrophs, indirect evidence suggests that non-cyanobacterial diazotrophs (NCDs) might also be important. One widely used approach for understanding diazotroph diversity and biogeography is polymerase chain reaction (PCR)-amplification of a portion of the *nifH* gene, which encodes a structural component of the N_2_-fixing enzyme complex, nitrogenase. An array of bioinformatic tools exists to process *nifH* amplicon data, however, the lack of standardized practices has hindered cross-study comparisons. This has led to a missed opportunity to more thoroughly assess diazotroph biogeography, diversity, and their potential contributions to the marine N cycle. To address these knowledge gaps a bioinformatic workflow was designed that standardizes the processing of *nifH* amplicon datasets originating from high-throughput sequencing (HTS). Multiple datasets are efficiently and consistently processed with a specialized DADA2 pipeline to identify amplicon sequence variants (ASVs). A series of customizable post-pipeline stages then detect and discard spurious *nifH* sequences and annotate the subsequent quality-filtered *nifH* ASVs using multiple reference databases and classification approaches. This newly developed workflow was used to reprocess nearly all publicly available *nifH* amplicon HTS datasets from marine studies, and to generate a comprehensive *nifH* ASV database containing 7909 ASVs aggregated from 21 studies that represent the diazotrophic populations in the global ocean. For each sample, the database includes physical and chemical metadata obtained from the Simons Collaborative Marine Atlas Project (CMAP). Here we demonstrate the utility of this database for revealing global biogeographical patterns of prominent diazotroph groups and highlight the influence of sea surface temperature. The workflow and *nifH* ASV database provide a robust framework for studying marine N_2_ fixation and diazotrophic diversity captured by *nifH* amplicon HTS. Future datasets that target understudied ocean regions can be added easily, and users can tune parameters and studies included for their specific focus. The workflow and database are available, respectively, in GitHub (https://github.com/jdmagasin/nifH-ASV-workflow) and Figshare (https://doi.org/10.6084/m9.figshare.23795943.v1).

## 1 Introduction

Dinitrogen (N_2_) fixation, the reduction of N_2_ into bioavailable NH_3_ is a source of new nitrogen (N) in the oceans and can support as much as 70% of new primary production in N-limited oligotrophic gyres (Jickells et al., 2017). Over millennia, N_2_ fixation may balance the loss of N from the marine system through denitrification and annamox (Zehr and Capone, 2020). N_2_ fixation was thought to be performed exclusively by prokaryotes, yet it was recently demonstrated that the marine haptophyte alga, *Braarudosphaera bigelowii*, contains a cyanobacterially-derived organelle specialized for N_2_ fixation (Coale et al., 2024). Noting this exception, microorganisms able to fix N_2_ (diazotrophs), are broadly characterized into two main groups, cyanobacterial diazotrophs (those phylogenetically related to cyanobacteria) and non-cyanobacterial diazotrophs (NCDs). Historically, cyanobacterial diazotrophs have been considered the most important contributors to marine N_2_ fixation (Villareal, 1994; Capone et al., 2005). NCDs, first detected by Zehr et al. (1998), have since been demonstrated to be ubiquitous in pelagic marine waters, and are generally thought to be putative chemoheterotrophs with a highly diverse lineage that includes the massive phylum Proteobacteria as well as Firmicutes, Actinobacteria, and Chloroflexi (Turk-Kubo et al., 2022). However, their contribution of fixed N and their role in the global ocean is not well-understood (Moisander et al., 2017).

Diazotrophs are often present at low abundances relative to other members of ocean microbiomes, which makes them challenging to study (Moisander et al., 2017; Benavides et al.). Distinctive pigments and morphologies that enable some cyanobacterial diazotrophs to be identified by microscopy are lacking in many diazotrophs (Carpenter and Capone, 1983; Carpenter and Foster, 2002), including NCDs. Furthermore, many marine diazotrophs are uncultivated, which has required the use of cultivation-independent approaches such as PCR and quantitative PCR (qPCR) (Luo et al., 2012; Shao and Luo, 2022; Turk-Kubo et al., 2022). The *nifH* gene encodes the identical subunits of the Fe protein of nitrogenase, the enzyme that catalyzes the N_2_ fixation reaction, and contains both highly conserved and variable regions enabling its use as a phylogenetic marker and as a proxy for N_2_-fixing potential in marine ecosystems globally (Gaby and Buckley, 2011).

Although the importance of marine N_2_ fixation is well-established, knowledge gaps remain, and discoveries continue to be made (Zehr and Capone, 2020). For example, high-throughput sequencing (HTS) of *nifH* amplicons is expanding our knowledge of diazotroph biogeography and activity and has revealed surprising new diversity. However, HTS studies often utilize different or custom software pipelines and parameters, rendering direct comparisons between studies difficult. Additionally, many studies do not address the full breadth of diazotrophic diversity because they focus on cyanobacterial diazotrophs while providing only a superficial analysis of the NCDs present. The resulting lack of information on NCD *in situ* distributions limits our understanding of diazotroph ecology and N_2_ fixation as well as our ability to predict how these populations will respond, e.g., trait-based ecological models, to a continually changing ocean.

To address these issues, we compiled published *nifH* amplicon HTS datasets along with two new datasets. Twenty-one studies were reprocessed by our newly developed software workflow, which streamlines the integration of multiple, large amplicon datasets for reproducible analyses. The workflow identifies amplicon sequence variants (ASVs) using a pipeline developed around DADA2 (Callahan et al., 2016) — the DADA2 *nifH* pipeline — and then executes rigorous post-pipeline stages to: remove spurious *nifH* ASVs; annotate the remaining quality-filtered ASVs using multiple reference databases and classification approaches; and obtain *in situ* and modeled environmental data for each sample from the Simons Collaborative Marine Atlas Project (CMAP; https://simonscmap.com). Although created to support research into N_2_ fixation (*nifH*), the complete workflow (ASV pipeline followed by the post-pipeline stages) can be adapted for use with other amplicon datasets, including other functional genes or taxonomic markers (16S rRNA genes), with some simple modifications.

In addition to the workflow, our efforts resulted in the construction of a comprehensive database of *nifH* ASVs with contextual metadata that will be a community resource for marine diazotroph investigations, enhancing comparability between previous and future *nifH* amplicon datasets. The *nifH* ASV database is available in Figshare (https://doi.org/10.6084/m9.figshare.23795943.v1). The entire workflow required to produce the *nifH* ASV database is available in two GitHub repositories, the DADA2 *nifH* pipeline (https://github.com/jdmagasin/nifH_amplicons_DADA2), and the post-pipeline stages (https://github.com/jdmagasin/nifH-ASV-workflow).

## 2 Data and Methods

### 2.1 Overview of *nifH* amplicon workflow and *nifH* ASV database generation

The full workflow is comprised of two parts: 1) the DADA2 *nifH* pipeline; and 2) a series of post-pipeline stages (Fig. 1). Required inputs for the pipeline are raw *nifH* amplicon sequencing reads and sample collection metadata (at minimum the latitude and longitude, depth and sample collection date and time) used to acquire environmental metadata from CMAP. Criteria for including publicly available datasets are detailed in Section 2.2.1.

**Figure 1:**
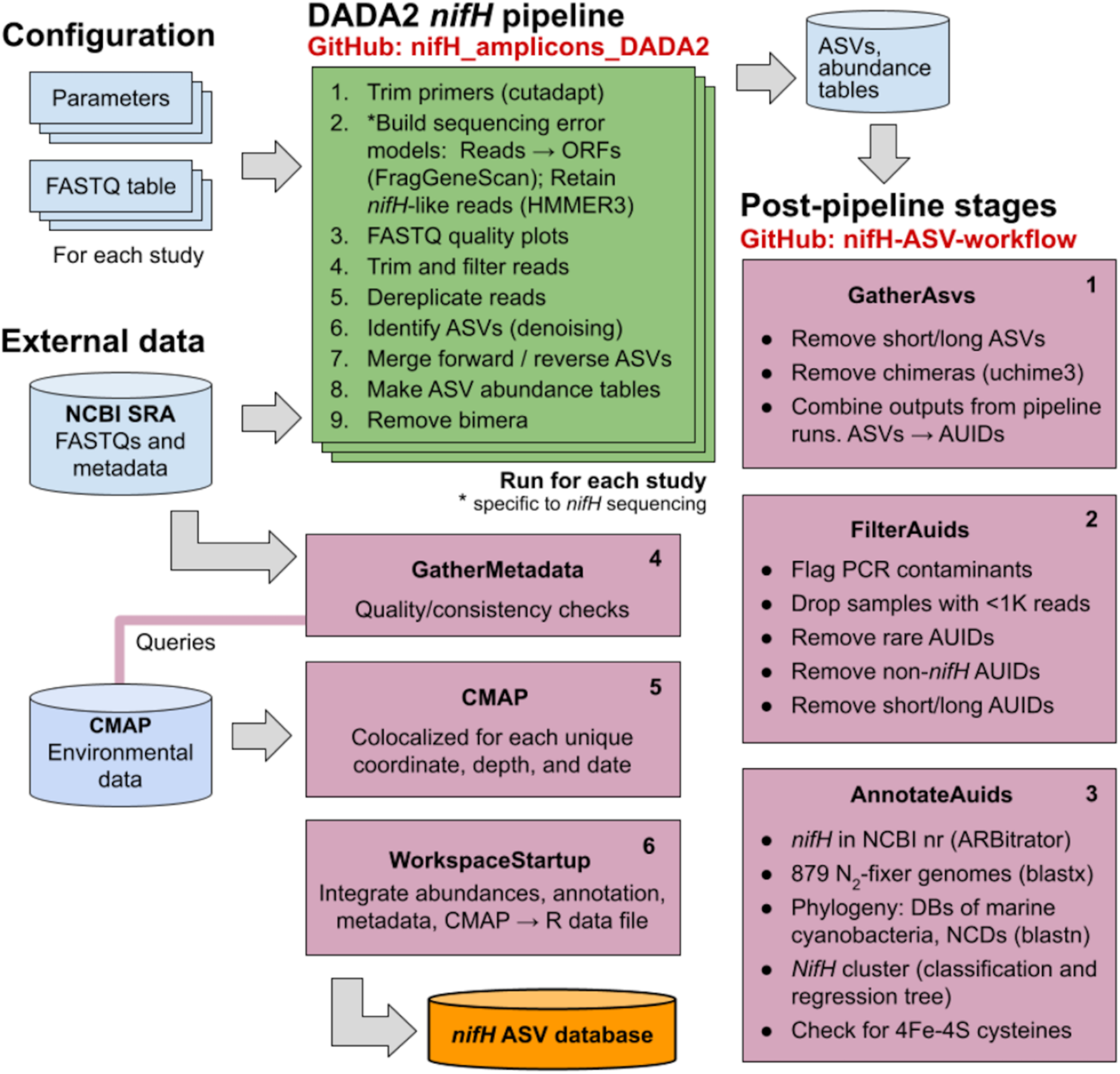
Schematic of the *nifH* amplicon data workflow. Data from all studies that met our criteria (Sect. 2.2) were downloaded from the NCBI Sequence Read Archive (SRA) and processed separately through the DADA2 *nifH* pipeline (green; Sect. 2.3.2), generally using identical parameters. ASV sequences and abundance tables from all studies were then combined and processed through each stage of the post-pipeline workflow (purple, Sect. 2.3.3) by executing the Makefile associated with each stage. Post-pipeline stages quality-filtered and then annotated the ASVs by reference to several *nifH* databases, and downloaded CMAP environmental data matched to the date, coordinates, and depth of each amplicon dataset. The main output of the entire workflow (pipeline and post-pipeline) is the *nifH* ASV database, which is available in Figshare (https://doi.org/10.6084/m9.figshare.23795943.v1). The workflow is maintained in two GitHub repositories, one for the DADA2 *nifH* pipeline (https://github.com/jdmagasin/nifH_amplicons_DADA2) and one for the post-pipeline stages (https://github.com/jdmagasin/nifH-ASV-workflow).

The DADA2 software package is frequently used for processing 16/18S rRNA gene amplicon sequencing data due to its ability to remove base calling errors (“denoising”) and thereby infer error-free ASVs (Callahan et al., 2016). We have developed a customizable pipeline to improve the error models utilized by DADA2 by training them only on reads in a dataset that are valid *nifH* sequences (not PCR artifacts). The DADA2 pipeline runs from the command line in a Unix-like shell, moving through nine steps (Fig. 1 DADA2 *nifH* pipeline) described in Section 2.3.2 for each study independently. After the DADA2 pipeline is completed, outputs from all studies are integrated and refined by the six post-pipeline stages of the workflow, which perform additional quality filtering (e.g., size- and abundance-based selection), identify and remove spurious sequences (e.g., potential contaminants and non-target sequences), and annotate the ASVs (Fig. 1 Post-pipeline stages). By considering ASVs from all studies simultaneously, the workflow considers rare ASVs that might be discarded as irrelevant in a single-study analysis. Workflow stages are executed manually by running their associated Makefiles and Snakefiles within a Unix-like shell.

The workflow generates the final data product published in this work, the *nifH* ASV database, which includes ASV sequences, abundance and annotation tables, sample collection metadata, and sample environmental data from CMAP (Fig. 1). The database is available in Figshare (https://doi.org/10.6084/m9.figshare.23795943.v1) as a set of tables (comma-separated value files) and an ASV FASTA file. However, these are also provided within an R data file, workspace.RData, in the WorkspaceStartup directory in the workflow GitHub repository, for users who wish to analyze, curate, or customize the database using R packages for ecological analysis. All documentation, scripts, and data needed to run the workflow and produce the *nifH* ASV database are provided in the workflow GitHub repository (https://github.com/jdmagasin/nifH-ASV-workflow). This includes pre-generated pipeline results for each of the 21 studies as well as the pipeline parameters files.

In summary, the workflow facilitates the systematic and reproducible exploration of *nifH*-based diversity within microbial communities and was applied to available *nifH* amplicon data to generate a globally distributed *nifH* ASV database. Together the workflow and *nifH* ASV database will serve as valuable community resources, fostering future investigations while ensuring comparability between previous and forthcoming studies. In the following sections, detailed descriptions of each stage of the workflow are provided.

### 2.2 Compilation of *nifH* amplicon studies

#### 2.2.1 Published studies

We compiled all publicly available *nifH* amplicon HTS data that were generated using the nifH1-4 primers (Zani, 1999; Zehr and Mcreynolds, 1989) and subsequently sequenced on the Illumina MiSeq/HiSeq platform totaling 19 studies (Table 1). Limiting the scope to investigations that used the same amplification primers enabled a more tractable comparison across studies by different research groups that employed varying approaches to sample collection and preparation for sequencing by different centers. Datasets were downloaded directly from the National Center for Biotechnology Information (NCBI) Sequencing Read Archive (SRA) using the GrabSeqs tool (Taylor et al., 2020) by specifying the study’s NCBI project accession. Each dataset obtained included paired-end sequencing reads (in FASTQ files) and a table with the collection metadata for each sample. Some datasets could not be retrieved directly from the SRA and were obtained directly from the authors (Supplementary Table 1). Note that we did not include studies where data was generated from experimental perturbations or particle enrichments (Supplementary Table 1). Data were last accessed from NCBI SRA on 17 April 2024.

**Table 1:**
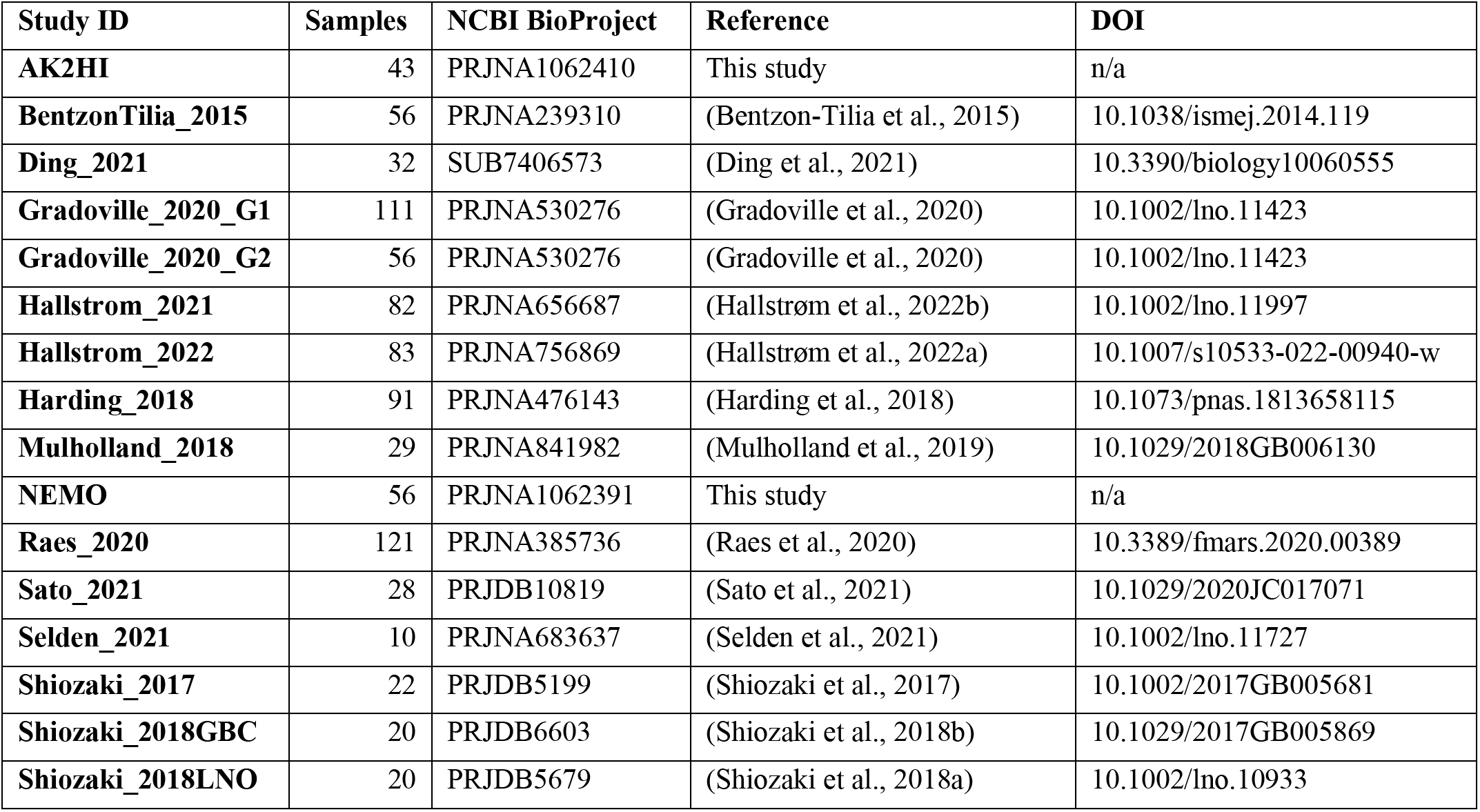
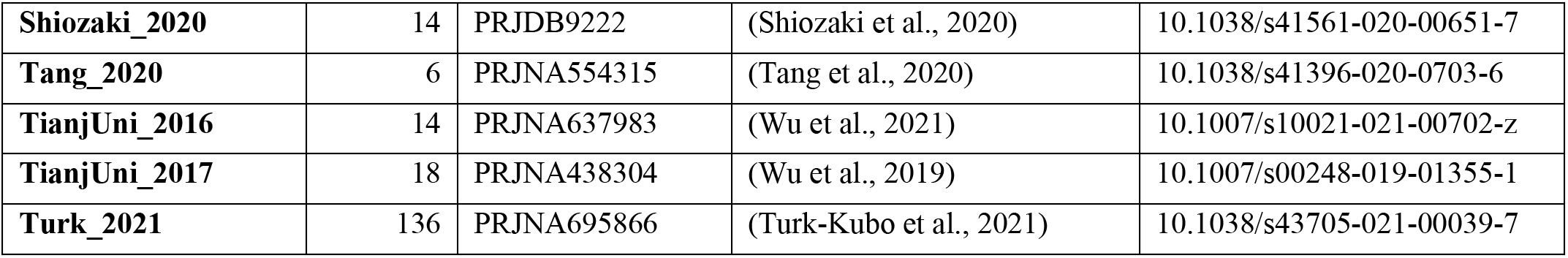
Information on the studies compiled to generate the *nifH* ASV database. All compiled studies and associated information. This includes the study ID used to refer to each dataset, the number of samples, NCBI BioProject accession, a reference to each publication and its corresponding DOI.

| Study ID | Samples | NCBI BioProject | Reference | DOI |
| --- | --- | --- | --- | --- |
| AK2HI | 43 | PRJNA1062410 | This study | n/a |
| BentzonTilia_2015 | 56 | PRJNA239310 | (Bentzon-Tilia et al., 2015) | 10.1038/ismej.2014.119 |
| Ding_2021 | 32 | SUB7406573 | (Ding et al., 2021) | 10.3390/biology10060555 |
| Gradoville_2020_G1 | 111 | PRJNA530276 | (Gradoville et al., 2020) | 10.1002/lno.11423 |
| Gradoville_2020_G2 | 56 | PRJNA530276 | (Gradoville et al., 2020) | 10.1002/lno.11423 |
| Hallstrom_2021 | 82 | PRJNA656687 | (Hallstrøm et al., 2022b) | 10.1002/lno.11997 |
| Hallstrom_2022 | 83 | PRJNA756869 | (Hallstrøm et al., 2022a) | 10.1007/s10533-022-00940-w |
| Harding_2018 | 91 | PRJNA476143 | (Harding et al., 2018) | 10.1073/pnas.1813658115 |
| Mulholland_2018 | 29 | PRJNA841982 | (Mulholland et al., 2019) | 10.1029/2018GB006130 |
| NEMO | 56 | PRJNA1062391 | This study | n/a |
| Raes_2020 | 121 | PRJNA385736 | (Raes et al., 2020) | 10.3389/fmars.2020.00389 |
| Sato_2021 | 28 | PRJDB10819 | (Sato et al., 2021) | 10.1029/2020JC017071 |
| Selden_2021 | 10 | PRJNA683637 | (Selden et al., 2021) | 10.1002/lno.11727 |
| Shiozaki_2017 | 22 | PRJDB5199 | (Shiozaki et al., 2017) | 10.1002/2017GB005681 |
| Shiozaki_2018GBC | 20 | PRJDB6603 | (Shiozaki et al., 2018b) | 10.1029/2017GB005869 |
| Shiozaki_2018LNO | 20 | PRJDB5679 | (Shiozaki et al., 2018a) | 10.1002/lno.10933 |
| **Shiozaki_2020** | 14 | PRJDB9222 | (Shiozaki et al., 2020) | 10.1038/s41561-020-00651-7 |
| **Tang_2020** | 6 | PRJNA554315 | (Tang et al., 2020) | 10.1038/s41396-020-0703-6 |
| **TianjUni_2016** | 14 | PRJNA637983 | (Wu et al., 2021) | 10.1007/s10021-021-00702-z |
| **TianjUni_2017** | 18 | PRJNA438304 | (Wu et al., 2019) | 10.1007/s00248-019-01355-1 |
| **Turk_2021** | 136 | PRJNA695866 | (Turk-Kubo et al., 2021) | 10.1038/s43705-021-00039-7 |

Sample quality was validated prior to processing through the DADA2 *nifH* pipeline. Samples were discarded if they did not contain unmerged pairs of forward and reverse reads with properly oriented primer sequences (Supplementary Table 1). There were two exceptions, studies by Shiozaki et al. (2017) and Shiozaki et al. (2018b), that used mixed-orientation sequence libraries and required preprocessing. The reads in each of these studies were partitioned by whether they captured the coding or template strand of *nifH*, determined by primer orientation. Because HTS sequence quality generally degrades from 5’ to 3’, the partitioned data were run separately through the pipeline to preserve their sequencing error profiles for DADA2. The ASVs from the misoriented reads (e.g. forward reads with template sequence) were then reverse-complemented and combined with the properly oriented ASVs into a single ASV abundance table and FASTA file. Table 1 and Supplementary Table 1 provide information for obtaining the raw FASTQ files for all samples evaluated for the *nifH* ASV database including information regarding studies excluded from the database.

#### 2.2.2 Unpublished *nifH* amplicon datasets

Additional *nifH* gene HTS datasets were included from DNA samples collected on two cruises in the North Pacific. One was a transect cruise across the Eastern North Pacific (NEMO; R/V New Horizon, August 2014; Shilova et al., 2017), and the other was a transect cruise from Alaska to Hawaii (AK2HI; R/V Kilo Moana, September 2017). Euphotic zone samples were collected from Niskin bottles deployed on a CTD-rosette (NEMO) or from the underway water system (5 m; AK2HI). NEMO samples (2-4 L) were filtered through 0.2 µm and 3 µm pore-size filters (in series), while AK2HI samples (ca. 2 L) were filtered through 0.2 µm pore-size filters using gentle peristaltic pumping. Filters were dried, flash frozen and stored at -80°C until processing. DNA was extracted using a modified DNeasy Plant Kit (Qiagen, Germantown, MD) protocol, described in detail in Moisander et al. (2008), with on-column washing steps automated by a QIAcube (Qiagen).

Partial *nifH* DNA sequences were PCR-amplified using the nifH1-4 primers in a nested *nifH* PCR assay (Zani, 1999; Zehr and Mcreynolds, 1989) according to details in Cabello et al. (2020). All samples were amplified in duplicate and pooled prior to sequencing. A targeted amplicon sequencing approach was used to create barcoded libraries as described in Green et al. (2015), using 5’ common sequence linkers (Moonsamy et al., 2013) on second round primers, nifH1 and nifH2. Sequence libraries were prepared at the DNA Service Facility at the University of Illinois at Chicago, and multiplexed amplicons were bidirectionally sequenced (2 × 300 bp) using the Illumina MiSeq platform at the W.M. Keck Center for Comparative and Functional Genomics at the University of Illinois at Urbana-Champaign. Samples were multiplexed to achieve ca. 40,000 high quality paired reads per sample. The AK2HI and NEMO datasets can be found in the SRA (BioProjects PRJNA1062410 and PRJNA1062391, respectively).

#### 2.2.3 Sample collection data and co-localized CMAP environmental data

Sample collection data (e.g. coordinates, depth, date) and environmental data provide essential context for the interpretation of diazotroph ‘omics datasets. Large-scale multivariate analyses depend on properly formatted, complete, and ideally quality checked metadata from consistently collected and analyzed measurements. However, accessibility to this information is often limited (especially environmental data) for datasets published across multiple decades. Therefore, we first obtained sample collection metadata from the SRA, and corrected or flagged errors and inconsistencies in the GatherMetadata stage of our post- pipeline workflow (described below), to ensure consistency and completeness. For each sample, the geographic coordinates, depth, and collection date (at local noon) from the SRA were used to query the Simons Collaborative Marine Atlas Project on 24 March 2023 (CMAP; https://simonscmap.com/; Ashkezari et al., 2021) for co-localized environmental data using a custom script (query_CMAP.py) in the CMAP stage of the workflow (Fig. 1). CMAP is an open-source data portal designed for retrieving, visualizing, and analyzing diverse ocean datasets including research cruise-based and autonomous measurements of biological, chemical, and physical properties, multi-decadal global satellite products, and output from global-scale biogeochemical models. For each sample a mixture of 102 satellite derived and modeled environmental variables from the CMAP repository were obtained. These, along with the SRA collection data, are included in our database. Aggregated metadata for all samples are summarized in Supplementary Table 2 but a detailed description of environmental metadata can be found at the CMAP website (https://simonscmap.com/catalog). Metadata are available in the *nifH* ASV database (metaTab.csv for sample metadata and cmapTab.csv for environmental data).

### 2.3 Automated workflow for processing datasets with the DADA2 *nifH* pipeline

#### 2.3.1 Installation of the DADA2 *nifH* pipeline and the post-pipeline workflow

The workflow (Fig. 1) comprises two software projects installed from separate GitHub repositories, nifH_amplicons_DADA2 which comprises the ASV pipeline and ancillary scripts, and nifH-ASV-workflow which integrates pipeline results for all datasets with annotation and CMAP environmental data to produce the data deliverable of the present work, the *nifH* ASV database. Installation requires cloning the nifH_amplicons_DADA2 repository (https://github.com/jdmagasin/nifH_amplicons_DADA2) to a local machine and then downloading several external software packages using miniconda3. Detailed installation instructions are available from the GitHub homepage, as well as a small tutorial to verify the installation on a small *nifH* amplicon dataset and introduce the two main pipeline commands (organizeFastqs.R and run_DADA2_pipeline.sh). Altogether the installation and example take 30–40 min.

After installing the ASV pipeline, installation of the nifH-ASV-workflow proceeds similarly: Clone the GitHub repository (https://github.com/jdmagasin/nifH-ASV-workflow) and then download a few additional packages with miniconda3 (∼10 min to complete). For each study, the nifH-ASV-workflow includes the pipeline outputs (ASVs and abundance tables) which were used to create the *nifH* ASV database. Pipeline parameters and FASTQ input tables for each study are also provided for users who instead wish to rerun the pipeline starting from FASTQs downloaded from the SRA. Because the nifH-ASV-workflow includes data and parameters specific to the studies used in this work, it has a separate GitHub repository from the pipeline. However, we emphasize that together they comprise the *nifH* amplicon workflow in Fig. 1.

Adding a new dataset to the workflow can be summarized in four steps: (1) Start a Unix-like shell that includes the required software (by “activating” a minconda3 environment called nifH_ASV_workflow). (2) Generate ASVs for the new dataset by running it through the pipeline, likely multiple times to tune parameters (Table 2). Output can be placed in the Data directory alongside other studies used in this work, and SRA metadata must be added to Data/StudyMetadata. (3) Include the new ASVs in the workflow by appending rows to the table GatherASVs/asvs.noChimera.fasta_table.tsv, which has file paths to all ASV abundance tables. (4) For each stage shown in Fig. 1, run the associated Makefile or Snakefile from the Unix-like shell by executing "make" or "snakemake -c1 --use-conda", respectively. Documentation resides within each Makefile or Snakefile. Input tables from the post-pipeline workflow also have embedded documentation.

**Table 2.**
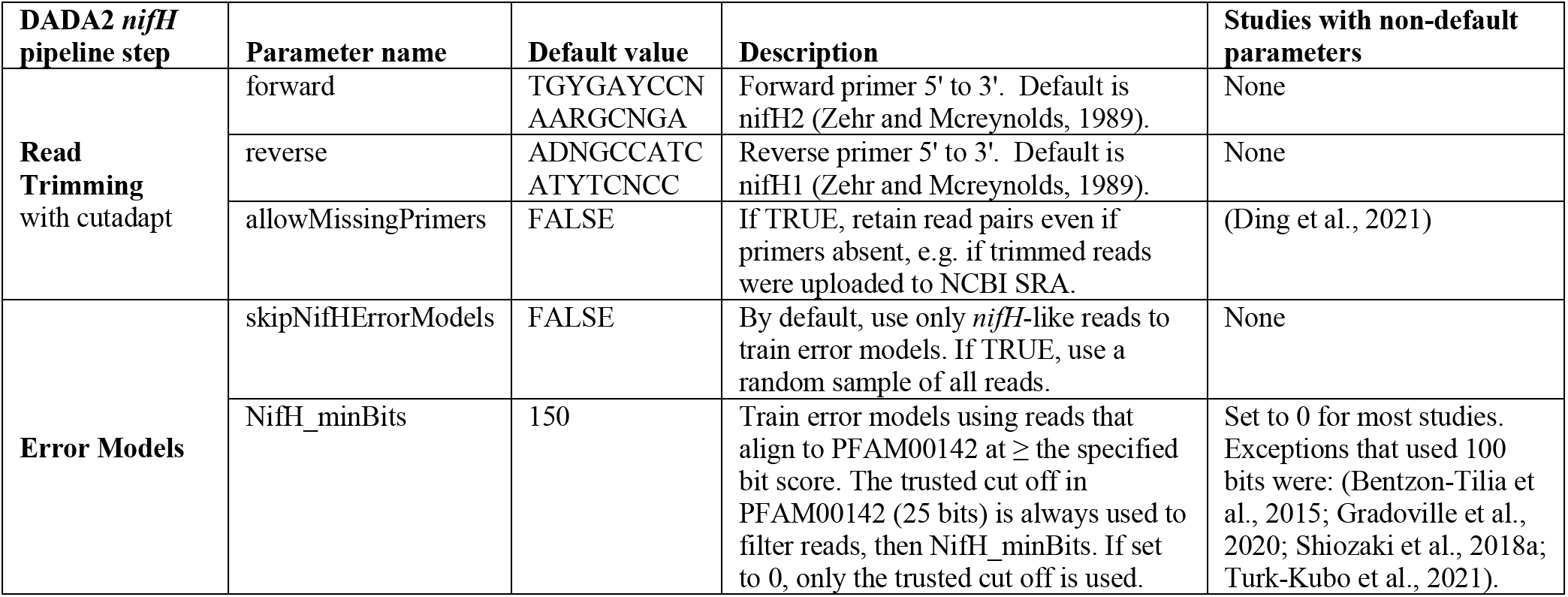
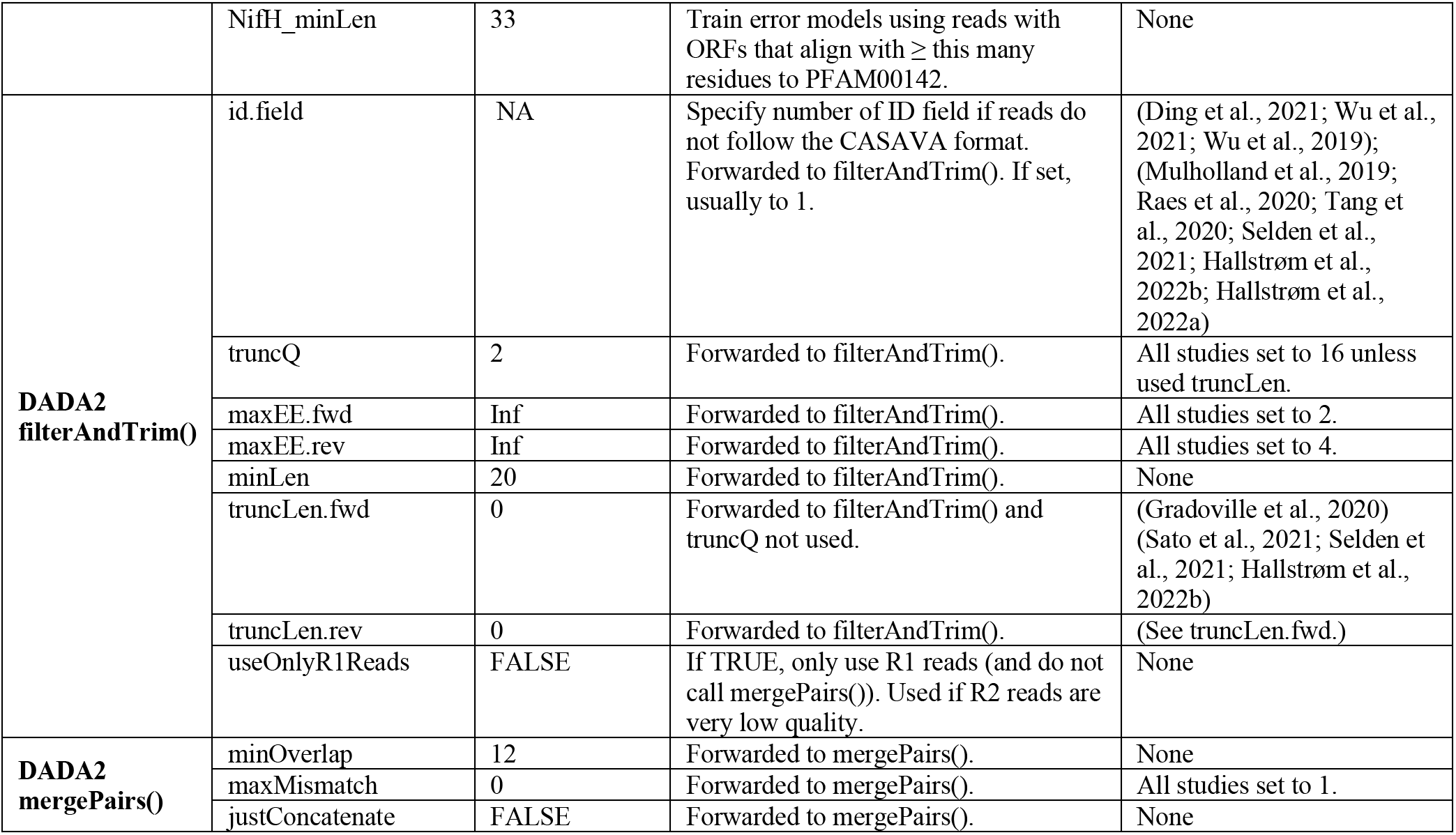
Parameters for controlling the DADA2 *nifH* pipeline. Default values can be overridden in the text file that is passed to run_DADA2_pipeline.sh. Parameters for "Read trimming" and "Error models" are used in steps 1 and 2 of the pipeline (Fig. 1). The remaining parameters are important for controlling how DADA2 trims and quality filters the reads, and merges forward and reverse sequences to create ASVs.

#### 2.3.2 DADA2 *nifH* pipeline

To encourage reproducible outputs and usage by non-programmers, the DADA2 pipeline (GitHub repository: nifH_amplicons_DADA2) is controlled by a plain text parameters file (Table 2) and a descriptive table of input samples (the “FASTQ map”). Since a study might include samples with vastly different diazotroph communities and relative abundances, potentially impacting ASV inferences by DADA2, the FASTQ map for a study enables samples to be partitioned into "processing groups” that are each run separately through DADA2. For example, in the present work processing groups usually partitioned the samples in a study by the unique combinations of collection station or date, nucleic acid type (DNA or RNA), size fraction, and collection depth. Pipeline outputs for each processing group are stored in a directory hierarchy with levels that follow the processing group definition. Partitioning datasets into processing groups greatly improves the overall speed of DADA2 and simplifies subsequent analyses that compare ASVs detected in different kinds of samples (e.g., detected versus transcriptionally active diazotrophs, or presence across different stations, depths, and/or size fractions). For generating the *nifH* ASV database, studies that met selection criteria (Sect. 2.2.1 and Table 1) were run through the pipeline using the study-specific FASTQ maps and parameters available in the Data directory of the nifH-ASV-workflow GitHub repository.

The DADA2 pipeline runs from the command line in a Unix-like shell, moving through 9 main steps (Fig. 1 DADA2 *nifH* pipeline): (1) trim reads of primers using cutadapt (Martin, 2011); (2) build sequencing error models; (3) make FASTQ quality plots; (4) trim and filter reads based on quality; (5) dereplicate; (6) denoise (ASV inference); (7) merge forward and reverse sequences; (8) make the ASV abundance table; and (9) remove bimera (Callahan et al., 2016 for steps 2 through 9). These steps will be familiar to DADA2 users, except that for step 2 the error models are trained only on *nifH*-like reads (discussed below). To run the pipeline on other functional genes, the parameters file would need to be edited to disable *nifH*-based error models and to include the expected primers. We again note that the DADA2 pipeline is distinct from the post-pipeline workflow stages which are specific to this work, but together they comprise the workflow in Fig. 1.

DADA2 parameters impact the ASV sequences identified, and the number of reads used. Thus, exploring parameters is essential for checking the robustness of ASVs (particularly rare ones) and their relative abundances. The DADA2 pipeline supports the optimization of parameters (Table 2). For example, one can trim each read based on its quality degradation (truncQ parameter to the DADA2 filterAndTrim function) or all reads at the same position determined by inspecting FASTQ quality plots. The pipeline allows one to rerun DADA2 steps 3-9, with outputs saved in separate, date-stamped directories. Read trimming and error models (steps 1-2) are unlikely to benefit much from parameter tuning, so the pipeline reuses outputs from those steps. Log files and diagnostic plots created by the pipeline are intended to facilitate parameter evaluation as well to capture statistics to support publication. Moreover, logs and other pipeline outputs are consistently formatted across pipeline runs, which enables scripts to aggregate and analyze results across datasets such as in our workflow.

Step 1 consisted only of read trimming using cutadapt (Martin, 2011). Raw reads were trimmed and retained only when read pairs for which the forward (nifH2) and reverse (nifH1) primers were both found on the R1 and R2 reads, respectively. DADA2 sequencing error models were built at step 2 using only the reads predicted to be *nifH*, rather than a subsample of all reads as in typical use of DADA2. Reads likely to encode *nifH* were identified as follows: FragGeneScan (version 1.31, (Rho et al., 2010)) was used to predict open reading frames (ORFs) on R1 reads which were then aligned to the nitrogenase PFAM model (PF00142.20) using HMMer3 (hmmsearch version 3.3.2; hmmer.org). ORFs with >33 residues and a bit score that exceeded the trusted cut-off encoded in the model (25.0 bits) were retained. Prefiltering the reads aims to reduce effects of PCR artifacts on the error models. For some studies this approach resulted in increases (∼3–10 %) in the total percentage of reads retained in ASVs, and fewer total ASVs, compared to using error models based on a subsample of all reads. Adapting the pipeline to a different marker gene would only require substituting an appropriate PFAM model, or disabling step 2 (by setting skipNifHErrorModels to TRUE; Table 2), which forces the pipeline to make error models by subsampling from all reads. At step 4, DADA2 filterAndTrim() truncated reads at the first base with PHRED score ≤16 and discarded read pairs that had excessive errors (>2 for R1 reads, >4 for R2 reads) or were <20 bp. The PHRED quality cut off, which corresponds to a 2.5 % base call error rate, was complemented by conservative parameters for merging sequences: At most 1 base pair was allowed to mismatch in the forward and reverse sequence overlap of minimally 12 bp (stage 7). Dereplicating (step 5) and denoising,

ASV calling (step 6), generating an abundance table (step 8), and bimera detection (step 9), were all performed with default DADA2 parameters. Data sets that passed pre-processing steps (Table 1) were run through the DADA2 pipeline using mostly identical parameters (Table 2).

#### 2.3.3 Post-pipeline stages

The workflow post-pipeline stages (GitHub repository: nifH-ASV-workflow) combine the pipeline outputs, conduct further quality control steps, co-locate the samples with environmental data from the CMAP data portal, and annotate the ASVs (Fig. 1 Post-pipeline stages). Key outputs from the post-pipeline are: a unified FASTA with all the unique ASVs detected across all the studies (i.e. all samples); tables of ASV total counts and relative abundances in all studies; multiple annotations for each ASV by comparison to several *nifH* reference databases; and CMAP environmental data for each sample. These outputs comprise the *nifH* ASV database, and are all available within an R image file (workspace.RData) generated by the workflow which is included in the nifH-ASV-workflow repository. Provision as an R image will make the outputs immediately accessible to many researchers who prefer R due to its extensive packages for ecological analysis. The *nifH* ASV database is also available on Figshare (https://doi.org/10.6084/m9.figshare.23795943.v1). The remainder of this section describes each of the post- pipeline stages.

The GatherAsvs stage aggregated ASV sequences and abundances across all DADA2 pipeline runs (i.e. from all samples and studies). First, ASVs were filtered based on length. Chimera sequences were then removed using UCHIME3 denovo (Edgar, 2016) via VSEARCH (Rognes et al., 2016). Chimera sequences were identified within each sample, but the final classification was based on majority vote (chimera or not) across the samples in the processing group. Second, the GatherAsvs stage combined the non-chimeric ASVs from all studies into a single abundance table and FASTA file. Since each study is run independently through the DADA2 pipeline, ASV identifiers are not consistent across studies. Therefore, each unique ASV sequence was renamed with a new unique identifier of the form AUID.*i*, where AUID stands for **A**SV **U**niversal **ID**entifier. The scripts used to rename the ASVs (assignAUIDs2ASVs.R) and to create the new abundance table (makeAUIDCountTable.R) are available at the nifH_amplicons_DADA2 GitHub repository (in scripts.ancillary/ASVs_to_AUIDs). The script assignAUIDs2ASVs.R optionally takes an AUID reference FASTA so that AUIDs can be preserved as new datasets are added to future versions of the *nifH* ASV database.

Both rare and potential non-*nifH* sequences were assessed on the unified AUID tables in the next stage, FilterAuids (Fig. 1). First, possible contaminants were identified by the Makefile invocation of check_nifH_contaminants.sh, provided as an ancillary script in the pipeline GitHub repository. In brief, check_nifH_contaminants.sh first translated all ASVs into amino acid sequences using FragGeneScan (Rho et al., 2010), which were then compared using *blastp* to 26 contaminants known from previous *nifH* amplicon studies (Zehr et al., 2003; Goto et al., 2005; Farnelid et al., 2009; Turk et al., 2011). ASVs that aligned at >96 % amino acid identity to known contaminants were flagged. Next FilterAuids removed samples with ≤1000 reads, and rare ASVs, defined as those that did not have at least one read in at least two samples or ≥1000 reads in one sample.

Next, the ancillary script, classifyNifH.sh, was employed to identify and remove non-*nifH*-like sequences. The script utilized *blastx* to search each ASV against ∼44 K positive and ∼15 K negative examples of NifH protein sequences that were found in NCBI GenBank by ARBitrator (run on April 28, 2020; Heller et al., 2014). ASVs were classified based on the relative quality of their best hits in the two databases, similar to the "superiority" check in ARBitrator. An ASV was classified as positive if the E-value of its best positive hit was ≥10 times smaller than the E-value for the best negative hit, and vice versa for negative classifications. ASVs failing to meet these criteria were classified as ‘uncertain’. The *blastx* searches used the same effective sizes for the two databases (-dbsize 1000000), so that E-values could be compared, and retained up to 10 hits (-max_target_seqs 10).

The FilterAuids stage of the workflow exclusively discarded ASVs with negative classifications. “Uncertain” ASVs were retained as potential *nifH* sequences not in GenBank. In the last stage, FilterAuids excluded ASVs with lengths that fell outside 281–359 nucleotides, a size range which in our experience encompasses the majority of valid *nifH* amplicon sequences generated by nested PCR with nifH1–4 primers.

For each AUID in the *nifH* ASV database, we provide taxonomical annotations using several different approaches, encompassed by the AnnotateAuids stage (Fig. 1) and accessible through ancillary scripts in the GitHub repository (in scripts.ancillary/Annotation). The script blastxGenome879.sh enables a protein level comparison via *blastx* against a database of 879 sequenced diazotroph genomes (“genome879”, https://www.jzehrlab.com/nifh). Here, the closest cultivated relative for each AUID was determined by smallest E-value among alignments with ≥50 % amino acid identity and ≥90 % query sequence coverage. Cautious interpretation is suggested because the reference DB is small and contains only cultivable taxa. Similarly, the top nucleotide match of each AUID was identified by E-value within alignments possessing ≥70 % nt identity and ≥90 % query sequence coverage obtained by *blastn* against a curated database of *nifH* sequences (July 2017, https://wwwzehr.pmc.ucsc.edu/nifH_Database_Public/) by executing the blastnARB2017.sh script. Additionally, *nifH* cluster annotations were assigned to each ASV using the classification and regression tree (CART) method of Frank et al. (2016). This approach was implemented as part of a custom tool that predicted ORFs for the ASVs with FragGeneScan, then performed a multiple sequence alignment on the ORFs, and then applied the CART classifier. The tool is available as the ancillary script assignNifHclustersToNuclSeqs.sh.

The Makefile created and searched against two "phylotype" databases, one containing 223 *nifH* sequences from prominent marine diazotrophs including NCDs (Turk-Kubo et al., 2022) and another with 44 UCYN-A *nifH* oligotype sequences (Turk-Kubo et al., 2017). These databases were searched using *blastn* with the effective database size of the ARB2017 database (- dbsize set to ∼29 million bases) to enable E-value comparisons across all three searches. For each ASV, we provide phylotype annotations based on the top hit by E-value if the alignment had ≥97 % nt identity and covered ≥70 % of the ASV. Finally, ORFs for all ASVs were searched for highly conserved residues which are thought to coordinate the 4Fe-4S cluster in NifH, specifically for paired cysteines shortly followed by AMP residues (described in Schlessman et al. 1998). This simple check, performed by the script check_CCAMP.R, was intended to complement the reference-based annotations above. Presence of cysteines and AMP could be used to retain ASVs that have no close reference. Absence could be used to flag ASVs that, despite high similarity to a reference sequence, might not represent functional *nifH* (e.g. due to frameshifts).

Since the annotation scripts provided multiple taxonomic identifications for most of the AUIDs, a primary taxonomic ID was assigned for each AUID using the script make_primary_taxon_id.py. If a phylotype annotation (e.g., Gamma A) was assigned, this became the primary taxonomic ID; otherwise, cultivated diazotrophs from genome879 were used (e.g., “*Pseudomonas stutzeri*”). Finally, when neither a phylotype nor a cultivated diazotroph could be determined, the *nifH* cluster (e.g. “unknown 1G”) was used. AUIDs without an assigned *nifH* cluster or taxonomic rank below domain were removed from the final *nifH* ASV database unless paired cysteines and AMP were detected. This final data filtration step occurred in the WorkspaceStartup stage described below.

The CMAP stage was managed by a Snakefile that called the script query_cmap.py to query the CMAP data portal for co- localized environmental data (Fig. 1). The script was passed the main output from the GatherMetadata stage, metadata.cmap.tsv, a table of the collection coordinates, dates at local noon, and depths from all the samples. GatherMetadata reported any samples with missing metadata and ensured standardized formats for the required query fields. Additionally, query_cmap.py validated fields prior to querying CMAP. It should be noted that the precision of values obtained from CMAP depend on floating point arithmetic, not the significant digits of the underlying measurement or model. Therefore, prior to an analysis requiring high precision for specific CMAP variables, it is recommended to consult the original producer of the data to determine the significant digits.

The last stage of the workflow, WorkspaceStartup, filtered out AUIDs that had no annotation and then generated the final *nifH* ASV database, which is comprised of AUID abundance tables (counts and relative), AUID annotations, sample metadata and corresponding environmental data. These data are provided as text files (.csv and FASTA) within a single compressed file (.tgz) that is available in Figshare (https://doi.org/10.6084/m9.figshare.23795943.v1) as well as within the workflow GitHub repository within an R image file (workspace.RData).

### 2.4 Diazotroph biogeography from DNA dataset of the *nifH* ASV database

The DNA dataset, a custom version of the *nifH* ASV database restricted to DNA samples (representing a majority of the database, only removing 94 samples), was created to showcase the utility of the workflow. Additional data reduction steps were conducted, averaging replicates and samples from the same location but different size fractions, to enable comparisons between different sampling methodologies.

## 3 Results and Discussion

### 3.1 Generation of the marine *nifH* ASV database

All publicly available marine *nifH* amplicon HTS data from studies that met our criteria, including two new studies, were compiled in the present investigation (see Sect. 2.2 and Supplementary Table 1). Altogether 982 samples from 21 studies, comprising a total of 87.7 million reads (Table 3), were processed through the entire workflow, i.e., the DADA2 *nifH* pipeline (Sect. 2.2.2) as well as the post-pipeline stages (Sect. 2.2.3). The *nifH* ASV database, i.e., the ASV sequences, abundances, and annotations, as well as sample collection and CMAP environmental data, was generated from the 865 samples, 7909 ASVs, and 34.4 million reads that were retained by this workflow (Figs. 1 and 2 and Table 3). To our knowledge it is the only global database for marine diazotrophs detected using *nifH* HTS amplicon sequencing, with comprehensive, standardized ancillary data (Fig. 2 and Supplementary Table 2).

**Figure 2:**
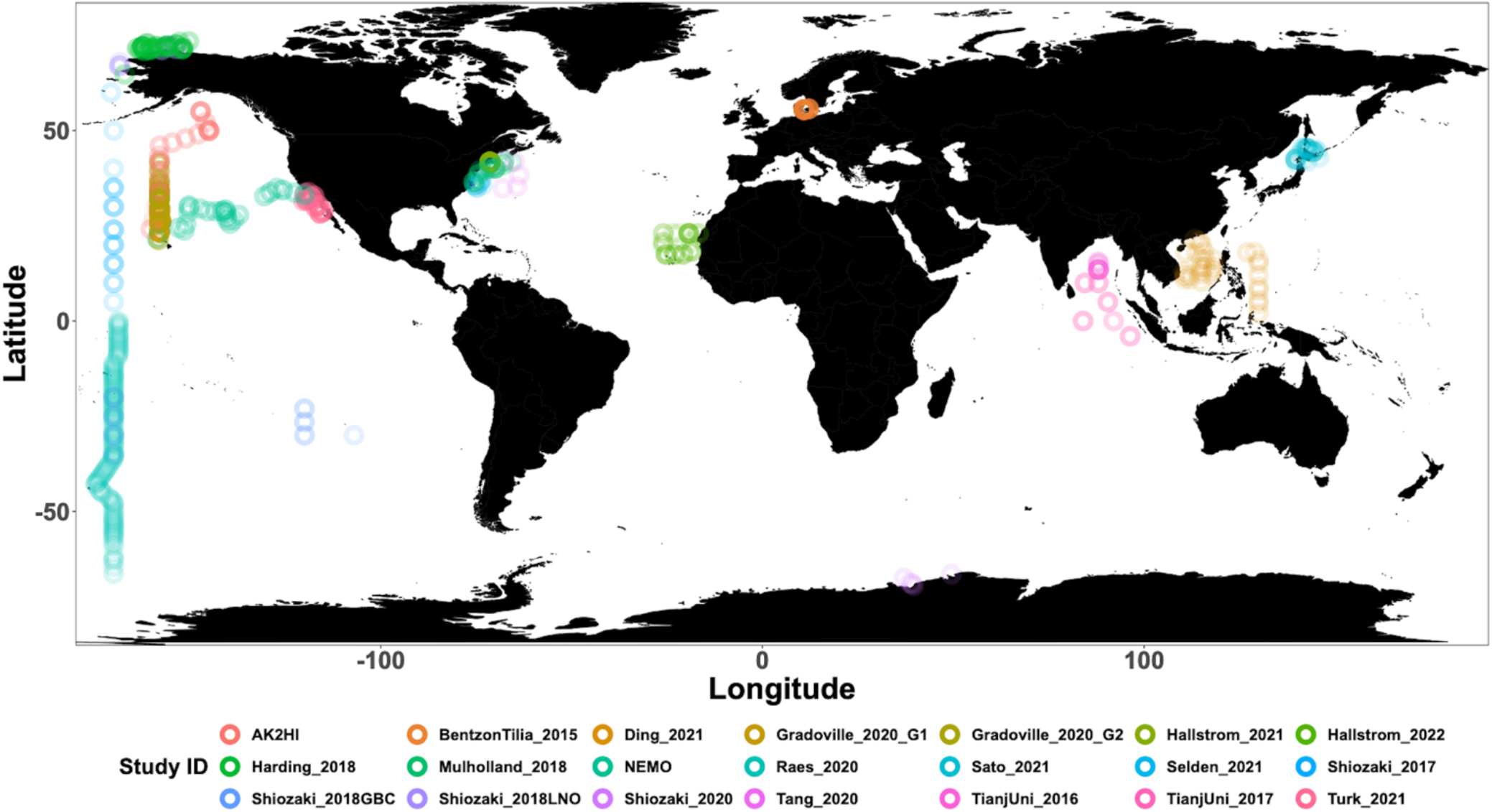
Global sampling distribution of the *nifH* ASV database. World map of sampling locations for the datasets compiled and processed to construct the *nifH* ASV database. See Table 1 for the citation source linked to each study ID.

**Table 3:** Summary of the full *nifH* workflow. The number of samples, ASVs, and reads retained through the entire workflow (the DADA2 *nifH* pipeline and major post-pipeline stages) to create the *nifH* ASV database. The vast majority ASVs that were removed by GatherAsvs fell outside 200–450 nt. WorkspaceStartup removed ASVs with no annotation and samples that had zero reads after ASV filtering.

|  | Initial | DADA2 pipeline | Gather Asvs | FilterAuids |  |  |  | Workspace Startup |
| --- | --- | --- | --- | --- | --- | --- | --- | --- |
|  |  |  |  | <1K reads in sample | rare | non-NifH | length |  |
| **Samples** | 982 | 982 | 982 | 894 | 890 | 890 | 890 | 865 |
| **ASVs** | n/a | 177,935 | 97,205 | 97,172 | 13,774 | 12,479 | 9,416 | 7,909 |
| **Reads (millions)** | 87.7 | 43.3 | 38.7 | 38.6 | 36.4 | 36.0 | 35.1 | 34.4 |

Interestingly, studies were affected differently by each step of the DADA2 *nifH* pipeline (Fig. 3 and Table 4). There were major losses of reads during ASV merging, with several studies retaining <25 % of their total reads by the end of the pipeline (i.e., BentzonTilia_2015, Hallstrom_2022, Shiozaki_2020, and TianjUni_2016), though on average about half the reads were retained across studies (Fig. 3 and Table 4).

**Figure 3:**
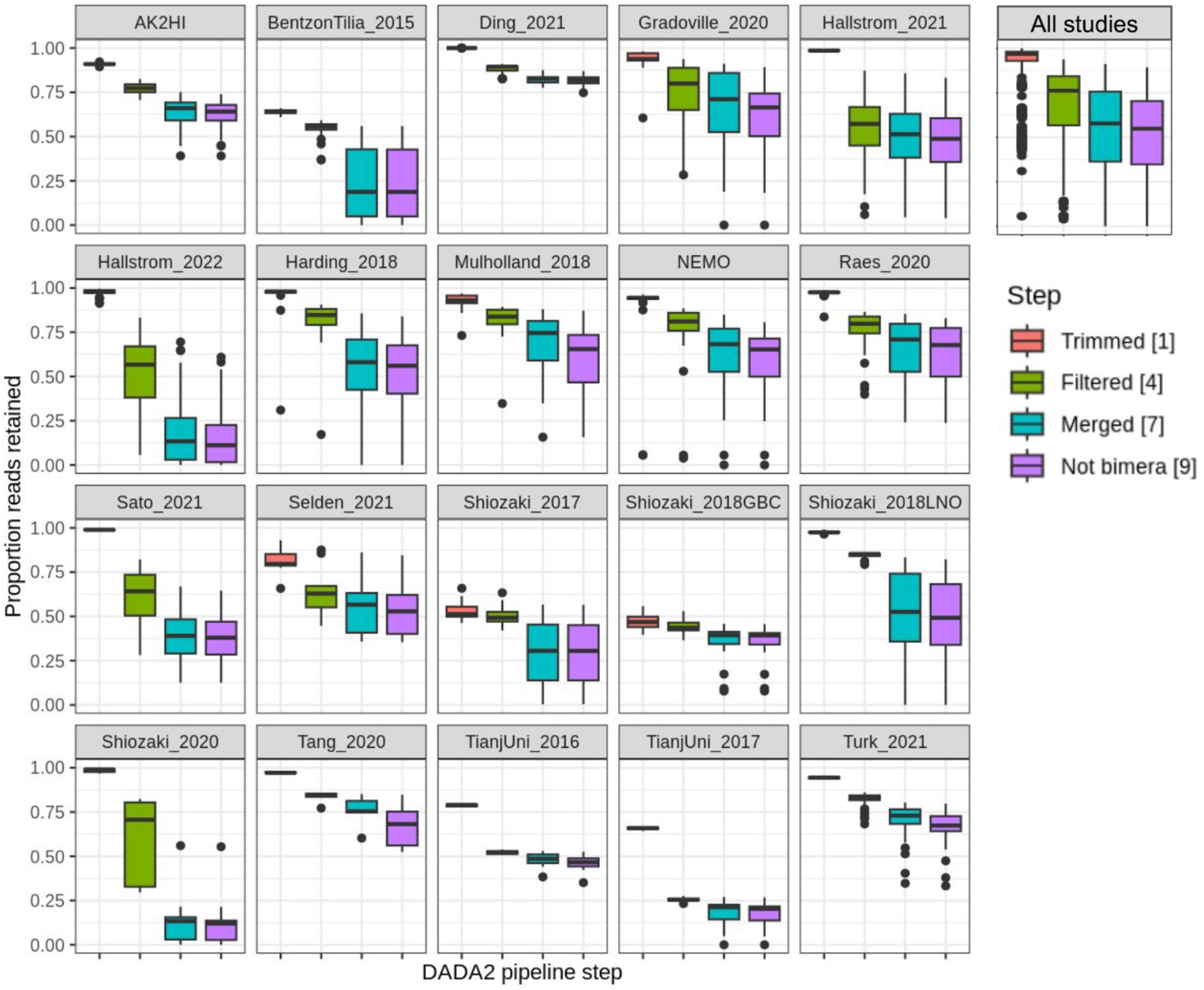
Study-specific retention of reads at each stage of the pipeline. The proportion of total reads in each sample that are retained at the completion of each step of the DADA2 *nifH* pipeline. Each box shows the distribution for samples in the indicated study (using Study IDs in Table 1), or for all samples together (top right). Proportions for Shiozaki_2017 and Shiozaki_2018GBC reflect that approximately half the amplicons were not in the orientation expected by the pipeline (see text). Numbers in the legend indicate pipeline steps in Fig. 1.

**Table 4:**
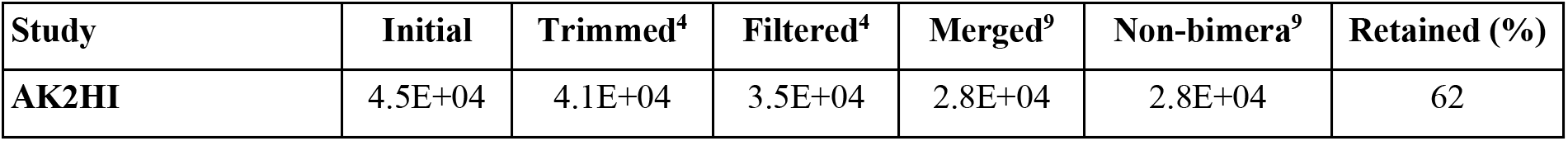
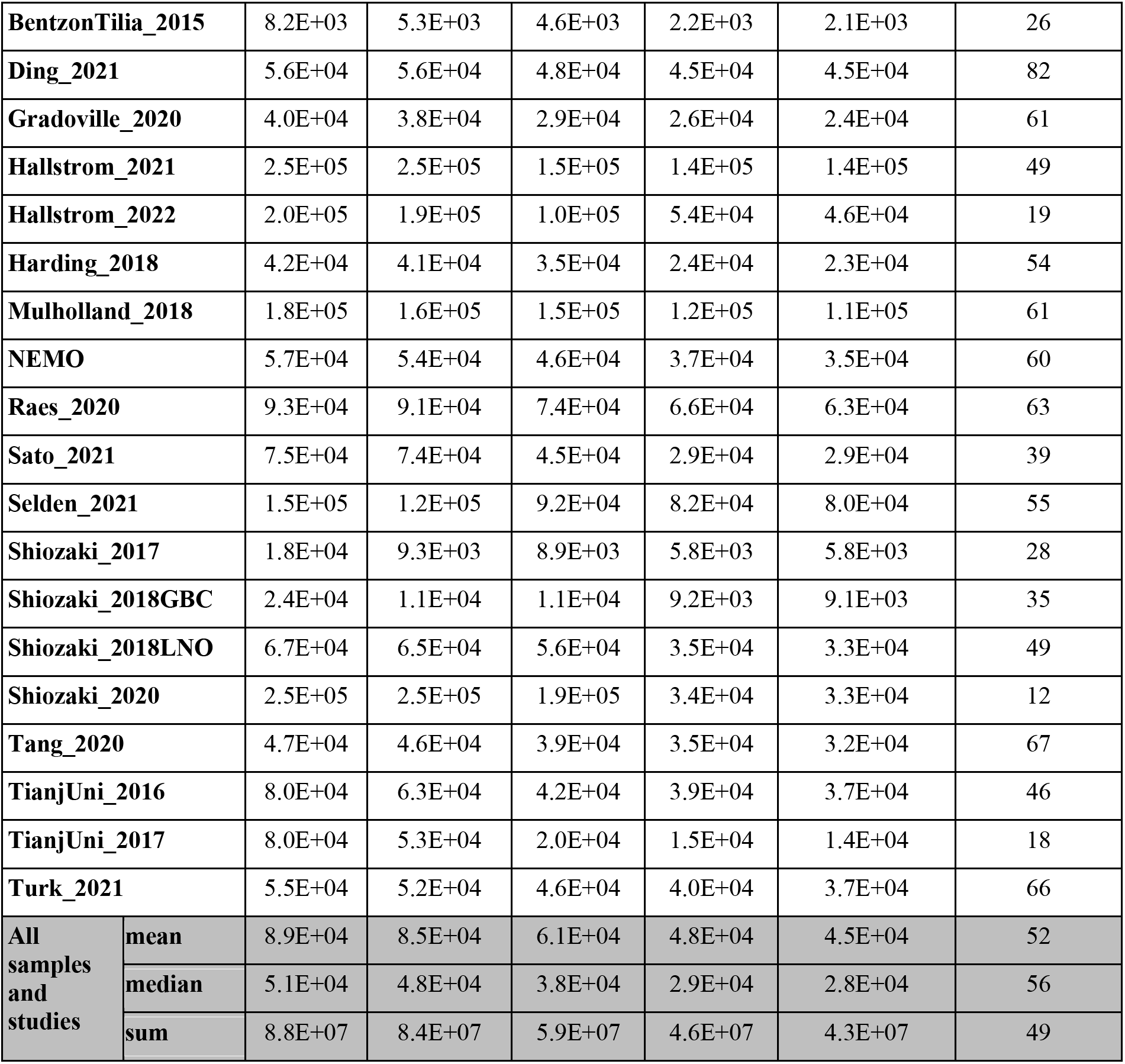
Quality filtering by the DADA2 *nifH* pipeline. For each study ID are shown the mean numbers of reads retained per sample at the end of each stage of the DADA2 *nifH* pipeline, as well as the mean percentage of reads retained. Statistics in the bottom three rows pool all samples. Initial, Trimmed^4^, Filtered^4^, and Merged^7^ and non-Bimera^9^ and their superscripts are specific to the pipeline steps in Fig. 1. At each step (column) the calculations include only the samples that have >0 reads.

| Study | Initial | Trimmed<sup>4</sup> | Filtered<sup>4</sup> | Merged<sup>9</sup> | Non-bimera<sup>9</sup> | Retained (%) |
| --- | --- | --- | --- | --- | --- | --- |
| AK2HI | 4.5E+04 | 4.1E+04 | 3.5E+04 | 2.8E+04 | 2.8E+04 | 62 |

|  |  |  |  |  |  |  |  |
| --- | --- | --- | --- | --- | --- | --- | --- |
| **BentzonTilia_2015** |  | 8.2E+03 | 5.3E+03 | 4.6E+03 | 2.2E+03 | 2.1E+03 | 26 |
| **Ding_2021** |  | 5.6E+04 | 5.6E+04 | 4.8E+04 | 4.5E+04 | 4.5E+04 | 82 |
| **Gradoville_2020** |  | 4.0E+04 | 3.8E+04 | 2.9E+04 | 2.6E+04 | 2.4E+04 | 61 |
| **Hallstrom_2021** |  | 2.5E+05 | 2.5E+05 | 1.5E+05 | 1.4E+05 | 1.4E+05 | 49 |
| **Hallstrom_2022** |  | 2.0E+05 | 1.9E+05 | 1.0E+05 | 5.4E+04 | 4.6E+04 | 19 |
| **Harding_2018** |  | 4.2E+04 | 4.1E+04 | 3.5E+04 | 2.4E+04 | 2.3E+04 | 54 |
| **Mulholland_2018** |  | 1.8E+05 | 1.6E+05 | 1.5E+05 | 1.2E+05 | 1.1E+05 | 61 |
| **NEMO** |  | 5.7E+04 | 5.4E+04 | 4.6E+04 | 3.7E+04 | 3.5E+04 | 60 |
| **Raes_2020** |  | 9.3E+04 | 9.1E+04 | 7.4E+04 | 6.6E+04 | 6.3E+04 | 63 |
| **Sato_2021** |  | 7.5E+04 | 7.4E+04 | 4.5E+04 | 2.9E+04 | 2.9E+04 | 39 |
| **Selden_2021** |  | 1.5E+05 | 1.2E+05 | 9.2E+04 | 8.2E+04 | 8.0E+04 | 55 |
| **Shiozaki_2017** |  | 1.8E+04 | 9.3E+03 | 8.9E+03 | 5.8E+03 | 5.8E+03 | 28 |
| **Shiozaki_2018GBC** |  | 2.4E+04 | 1.1E+04 | 1.1E+04 | 9.2E+03 | 9.1E+03 | 35 |
| **Shiozaki_2018LNO** |  | 6.7E+04 | 6.5E+04 | 5.6E+04 | 3.5E+04 | 3.3E+04 | 49 |
| **Shiozaki_2020** |  | 2.5E+05 | 2.5E+05 | 1.9E+05 | 3.4E+04 | 3.3E+04 | 12 |
| **Tang_2020** |  | 4.7E+04 | 4.6E+04 | 3.9E+04 | 3.5E+04 | 3.2E+04 | 67 |
| **TianjUni_2016** |  | 8.0E+04 | 6.3E+04 | 4.2E+04 | 3.9E+04 | 3.7E+04 | 46 |
| **TianjUni_2017** |  | 8.0E+04 | 5.3E+04 | 2.0E+04 | 1.5E+04 | 1.4E+04 | 18 |
| **Turk_2021** |  | 5.5E+04 | 5.2E+04 | 4.6E+04 | 4.0E+04 | 3.7E+04 | 66 |
| **All samples and studies** | **mean** | 8.9E+04 | 8.5E+04 | 6.1E+04 | 4.8E+04 | 4.5E+04 | 52 |
|  | **median** | 5.1E+04 | 4.8E+04 | 3.8E+04 | 2.9E+04 | 2.8E+04 | 56 |
|  | **sum** | 8.8E+07 | 8.4E+07 | 5.9E+07 | 4.6E+07 | 4.3E+07 | 49 |

Switching the trimming approach from one based on individual read quality profiles (using truncQ in Table 3) to fixed-length trimming based on overall quality profiles of the forward and reverse reads (using truncLen.fwd and truncLen.rev in Table 2) resulted in more reads being retained for some studies (Sato et al., 2021; Selden et al., 2021; Hallstrøm et al., 2022b; Gradoville et al., 2020). However, fixed-length trimming would have required the selection of trim lengths based on visual, qualitative assessments of hundreds of FASTQ quality plots which is difficult to accomplish in a systematic manner. For consistency we preferred to use nearly identical parameters for most studies (Table 3).

Post-pipeline stages of the workflow further refined the data (detailed in Methods) (Fig. 4). First, GatherAsvs identified and removed 112 chimeras using uchime3 denovo (distinct from the bimera filtering done by the pipeline), and then removed 81 K ASVs that were far outside expected *nifH* lengths (200–450 nt). AUIDs were assigned to the remaining 97 K unique non- chimeric ASVs (comprising 38.7 million total reads; Tables 3 and 5). The GatherAsvs length filter had by far the largest impact of any post-pipeline quality filtering, removing 10 % of the reads from the pipeline. Next, FilterAuids dropped four poorly sequenced samples (7 K total reads), as they would likely misrepresent their diazotrophic communities, and then removed 83 K rare ASVs (2.3 million reads; Tables 3 and 5).

**Figure 4:**
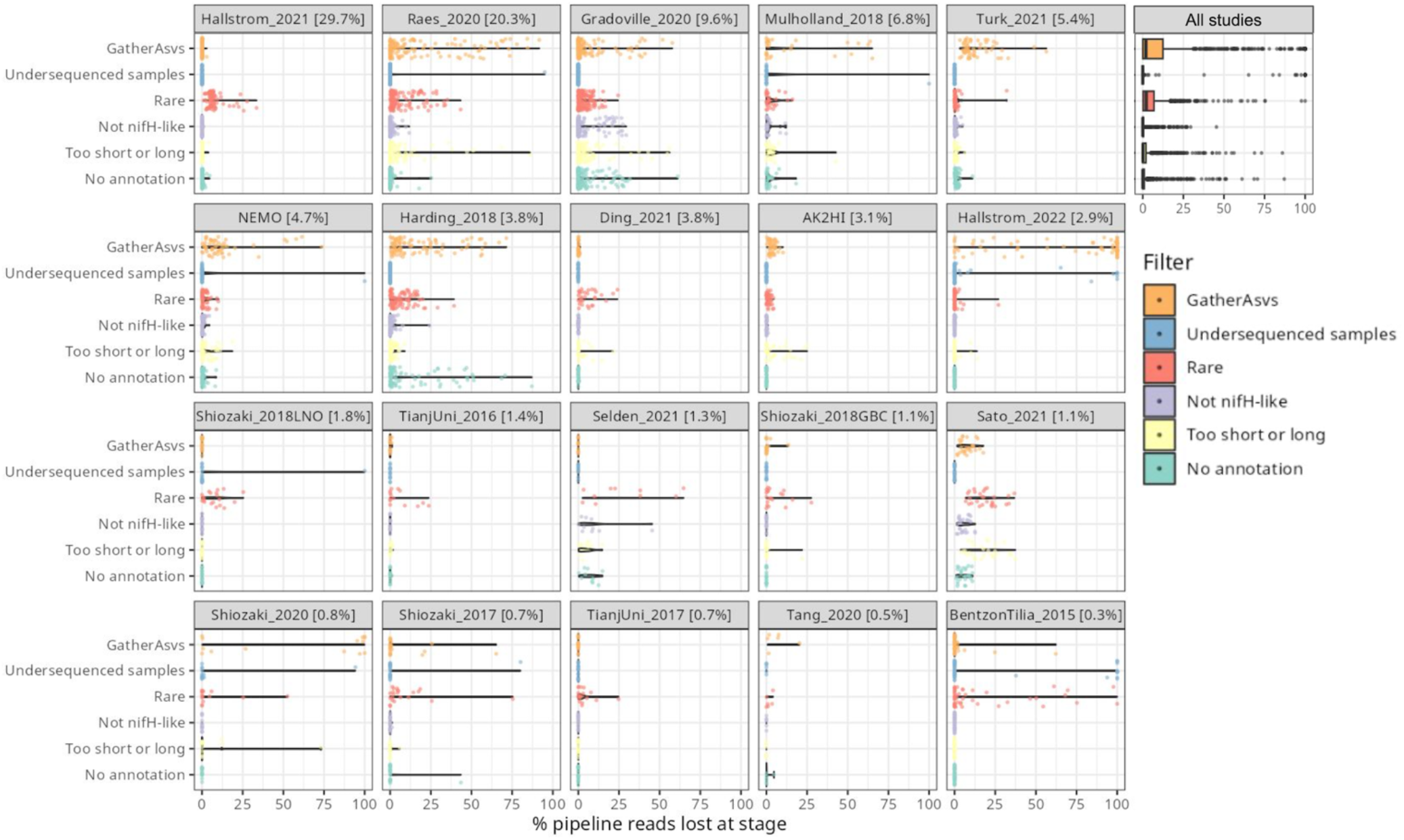
Study-specific retention of reads at each stage of the post-pipeline workflow. For each study the violin plots show how many reads from the pipeline were removed by GatherAsvs due to length, the four filtering steps of FilterAuids, or WorkspaceStartup due to the ASV having no annotation (shown in Fig. 1). Losses for all samples combined are shown in the box plot (top right). Studies are ordered by contribution to the *nifH* ASV database, e.g. 29.7 % of all the reads in the database were from Hallstrom_2021.

**Table 5.** Quality filtering by the post-pipeline workflow. For each study are shown the mean numbers of reads per sample that were output by the DADA2 *nifH* pipeline and retained by the GatherAsvs, FilterAuids, and WorkspaceStartup stages of the post-pipeline workflow. The Retained (%) column has the mean percentages of reads retained per sample (relative to column DADA2 pipeline values). Additionally, the last three rows show the overall means, medians, and sums of reads across all samples and studies. Superscripts correspond to stage numbers in Fig. 1 Post-pipeline stages. The GatherAsvs^1^ column mainly reflects length filtering (200–450 nt), and the WorkspaceStartup^6^ column reflects discarding of ASVs that had no annotation. At each stage (column) the calculations include only the samples that have >0 reads.

| Study ID |  | DADA2<br>pipeline | Gather<br>Asvs<sup>1</sup> | FilterAuids<sup>2</sup> |  |  | Workspace<br>Startup<sup>6</sup> | Retained<br>(%) |
| --- | --- | --- | --- | --- | --- | --- | --- | --- |
|  |  |  |  | Rare | Non-NifH | Length |  |  |
| AK2HI |  | 2.8E+04 | 2.7E+04 | 2.7E+04 | 2.7E+04 | 2.5E+04 | 2.5E+04 | 90 |
| BentzonTilia_2015 |  | 2.1E+03 | 2.1E+03 | 2.6E+03 | 2.6E+03 | 2.6E+03 | 2.6E+03 | 85 |
| Ding_2021 |  | 4.5E+04 | 4.5E+04 | 4.2E+04 | 4.2E+04 | 4.1E+04 | 4.1E+04 | 91 |
| Gradoville_2020 |  | 2.4E+04 | 2.3E+04 | 2.2E+04 | 2.1E+04 | 2.1E+04 | 2.0E+04 | 80 |
| Hallstrom_2021 |  | 1.4E+05 | 1.4E+05 | 1.3E+05 | 1.3E+05 | 1.2E+05 | 1.2E+05 | 92 |
| Hallstrom_2022 |  | 4.6E+04 | 2.6E+04 | 3.8E+04 | 3.8E+04 | 3.4E+04 | 3.4E+04 | 50 |
| Harding_2018 |  | 2.3E+04 | 1.9E+04 | 1.8E+04 | 1.7E+04 | 1.7E+04 | 1.5E+04 | 64 |
| Mulholland_2018 |  | 1.1E+05 | 9.3E+04 | 9.3E+04 | 9.1E+04 | 8.7E+04 | 8.4E+04 | 72 |
| NEMO |  | 3.5E+04 | 3.1E+04 | 3.1E+04 | 3.1E+04 | 3.0E+04 | 3.0E+04 | 80 |
| Raes_2020 |  | 6.3E+04 | 5.8E+04 | 5.6E+04 | 5.6E+04 | 5.6E+04 | 6.0E+04 | 76 |
| Sato_2021 |  | 2.9E+04 | 2.7E+04 | 2.1E+04 | 2.0E+04 | 1.5E+04 | 1.4E+04 | 43 |
| Selden_2021 |  | 8.0E+04 | 8.0E+04 | 6.0E+04 | 5.2E+04 | 4.9E+04 | 4.4E+04 | 52 |
| Shiozaki_2017 |  | 1.2E+04 | 1.2E+04 | 1.1E+04 | 1.1E+04 | 1.1E+04 | 1.1E+04 | 83 |
| Shiozaki_2018GBC |  | 2.0E+04 | 2.0E+04 | 2.0E+04 | 2.0E+04 | 2.0E+04 | 2.0E+04 | 93 |
| Shiozaki_2018LNO |  | 3.3E+04 | 3.3E+04 | 3.3E+04 | 3.3E+04 | 3.3E+04 | 3.3E+04 | 92 |
| Shiozaki_2020 |  | 3.3E+04 | 2.8E+04 | 4.2E+04 | 4.2E+04 | 5.7E+04 | 5.7E+04 | 61 |
| Tang_2020 |  | 3.2E+04 | 3.0E+04 | 2.9E+04 | 2.9E+04 | 2.9E+04 | 2.9E+04 | 91 |
| TianjUni_2016 |  | 3.7E+04 | 3.7E+04 | 3.5E+04 | 3.5E+04 | 3.5E+04 | 3.5E+04 | 93 |
| TianjUni_2017 |  | 1.4E+04 | 1.4E+04 | 1.4E+04 | 1.4E+04 | 1.4E+04 | 1.4E+04 | 96 |
| Turk_2021 |  | 3.7E+04 | 3.3E+04 | 3.3E+04 | 3.2E+04 | 3.2E+04 | 3.2E+04 | 83 |
| All<br>samples<br>and<br>studies | mean | 4.5E+04 | 4.2E+04 | 4.1E+04 | 4.0E+04 | 4.0E+04 | 4.0E+04 | 79 |
|  | median | 2.8E+04 | 2.6E+04 | 2.6E+04 | 2.6E+04 | 2.5E+04 | 2.6E+04 | 90 |
|  | sum | 4.3E+07 | 3.9E+07 | 3.6E+07 | 3.6E+07 | 3.5E+07 | 3.4E+07 | 79 |

Finally, ASVs were removed if they were classified as non-*nifH*, based on a strong alignment to sequences in NCBI nr that ARBitrator (Heller et al., 2014) classified as non-*nifH*. Specifically, an ASV was classified as non-*nifH* if the ratio of E-values for its best negative and positive hits, among sequences classified by ARBitrator, was >10. A total of 96,095 of the 97,205 non-chimera ASVs had database hits which resulted in 40,448 positive, 12,977 negative, and 42,670 uncertain classifications. This approach was used to leverage ARBitrator’s high specificity for detecting *nifH* as well as to enable users to identify ASVs that have high percent identity matches to sequences in GenBank. An alternative approach would have been to classify the ASVs based on their alignments to HMMs for NifH versus NifH-like proteins (e.g. protochlorophyllide reductase), used by the NifMAP pipeline for *nifH* operational taxonomic units (Angel et al., 2018). Finally, FilterAuids removed ASVs with lengths outside 281–359 nt, a total of 974 K reads and 3063 ASVs (Figs. 1, 4 and Tables 3 and 5). After FilterAUIDs, the total number of samples in the dataset was reduced from 982 to 890 and the number of ASVs from 97,205 to 9416.

FilterAuids also flagged a total of 2000 ASVs as possible PCR contaminants. Although we opted to flag, not remove, these ASVs, the workflow can be easily altered to remove contaminants. Most studies contained low levels of contamination (≤1 %) based on our criteria. However, several studies were flagged with ∼9–30 % of their reads being similar to known contaminants. Identifying potential contaminants is challenging given their numerous sources, study specific nature (Zehr et al., 2003), and lack of control sequence data from blanks.

Next, AnnotateAuids assigned annotations using our three *nifH* reference databases and CART (Fig. 1). In total 7931 of the 9416 quality filtered ASVs were annotated, usually with multiple references (Supplementary Fig. 1). Most (7926 ASVs) had hits to both genome879 and ARB2017, likely because the 879 sequenced diazotrophs had *nifH* homologs in GenBank that were found by ARBitrator. Fewer ASVs had hits to the databases that targeted UCYN-A oligos (102 ASVs) and other marine diazotrophs (645 ASVs; 96 ASVs also had UCYN-A hits). Most ASVs (7905 total) were assigned to NifH clusters 1–4 by CART (respectively, 4100; 79; 3607; and 109 ASVs), including five ASVs that had no hits to our databases. The majority of ASVs (7749 total) had open reading frames (ORFs) that contained paired cysteines and AMP which might coordinate the 4Fe- 4S cluster, and all 7749 also had annotations from the reference databases or CART. A few ASVs had annotations but lacked residues to coordinate 4Fe-4S: 23 ORFs lacked the paired cysteines and another 159 ORFs had paired cysteines but not AMP, usually due to a substitution for M. The last step of AnnotateAuids assigned primary IDs (described above) to 7908 ASVs. In the final stage of the post-pipeline workflow, WorkspaceStartup retained these 7908 ASVs. One ASV, which had no phylogroup but did have paired cysteines and AMP, was also retained. In total the *nifH* ASV database had 7909 ASVs comprising 34.4 million reads (Table 3).

In the CMAP stage, sample collection metadata (date, latitude, longitude, and depth) were used to download CMAP environmental data (102 variables) for each sample in the *nifH* ASV database (Fig. 1). The CMAP data will enable analyses of potential factors that influence the global distribution of the diazotrophic community. Aggregated metadata for all samples are available in the *nifH* ASV database (metaTab.csv for sample metadata and cmapTab.csv for environmental data).

The last stage of the post-pipeline workflow is WorkspaceStartup, which generates the *nifH* ASV database (Fig. 1). ASVs with no annotation are removed as well as samples with zero total reads due to ASV filtering steps. The *nifH* ASV database consisted of 21 studies, 865 samples, 7909 AVS and 34.4 million total reads (Tables 3 and 5). The database is heavily biased toward euphotic zone DNA samples, with euphotic heuristically defined as follows: Samples were classified as coastal (< 200 km from a major landmass) or open ocean. Euphotic samples were then identified as those collected above a depth cut off, 50 m for coastal samples and 100 m for open ocean. Samples obtained from DNA (n=768) far exceeded those from RNA (n=94) extracts. Likewise, a majority of the samples were from the euphotic zone (789 compared to 73 from the aphotic zone). The database also includes replicate samples (n=256) and size fractionated samples (n=142).

### 3.2 Global *nifH* ASV database

#### 3.2.1. Sample Distribution

Investigations of N_2_ fixation and diazotrophic communities have focused on specific ocean regions and this is reflected by the uneven global distribution of *nifH* amplicon datasets in the *nifH* ASV database (Figs. 2, 5a, and 5b). There is an outsized influence of the northern hemisphere, especially in the Pacific Ocean where most of the database samples were located (429) and 69.7 % of these samples originated from the northern hemisphere (Figs. 2, 5a, 5b, and 6). Ten studies are found within the Pacific, with several containing >50 samples (Figs. 2 and 6). Notably, Raes_2020 (Raes et al., 2020) is the largest dataset stretching from the equator to the Southern Ocean, making up almost the entirety of the southern hemisphere Pacific samples (Figs. 2 and 6). Two new studies carried out in the North Pacific constitute the only previously unpublished data of the *nifH* ASV database (Table 1). AK2HI was a latitudinal transect from Alaska (U.S.) to Hawaii (U.S.) and NEMO was a longitudinal transect across the Eastern North Pacific from San Diego, CA (U.S.) to Hawaii (U.S.) (Fig. 2; Sect. 2.2.2). The amplicon data compiled for the *nifH* ASV database was primarily generated from DNA, with most RNA samples deriving from Atlantic Ocean studies and no contribution from RNA samples in the Arctic or Indian Oceans (Fig. 6).

**Figure 5.**
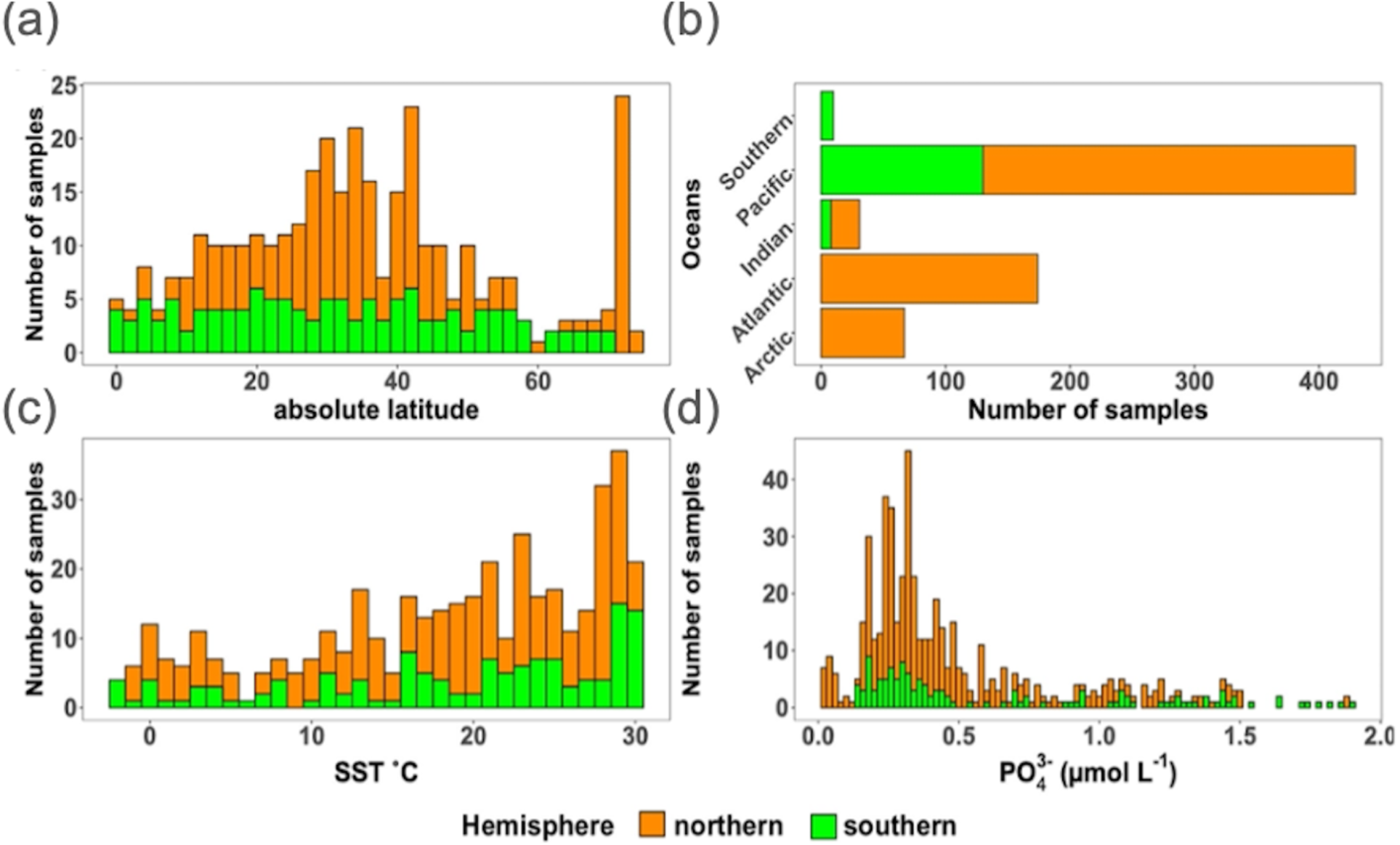
Location, temperature, and phosphate distributions of the *nifH* ASV database. The number of samples from the *nifH* ASV database by (a) absolute latitude, (b) the world’s oceans, (c) sea surface temperature (SST, °C) and (d) Pisces-derived PO_43-_ (µmol L^-1^). Environmental data, (c) and (d), were retrieved from the CMAP data portal.

**Figure 6.**
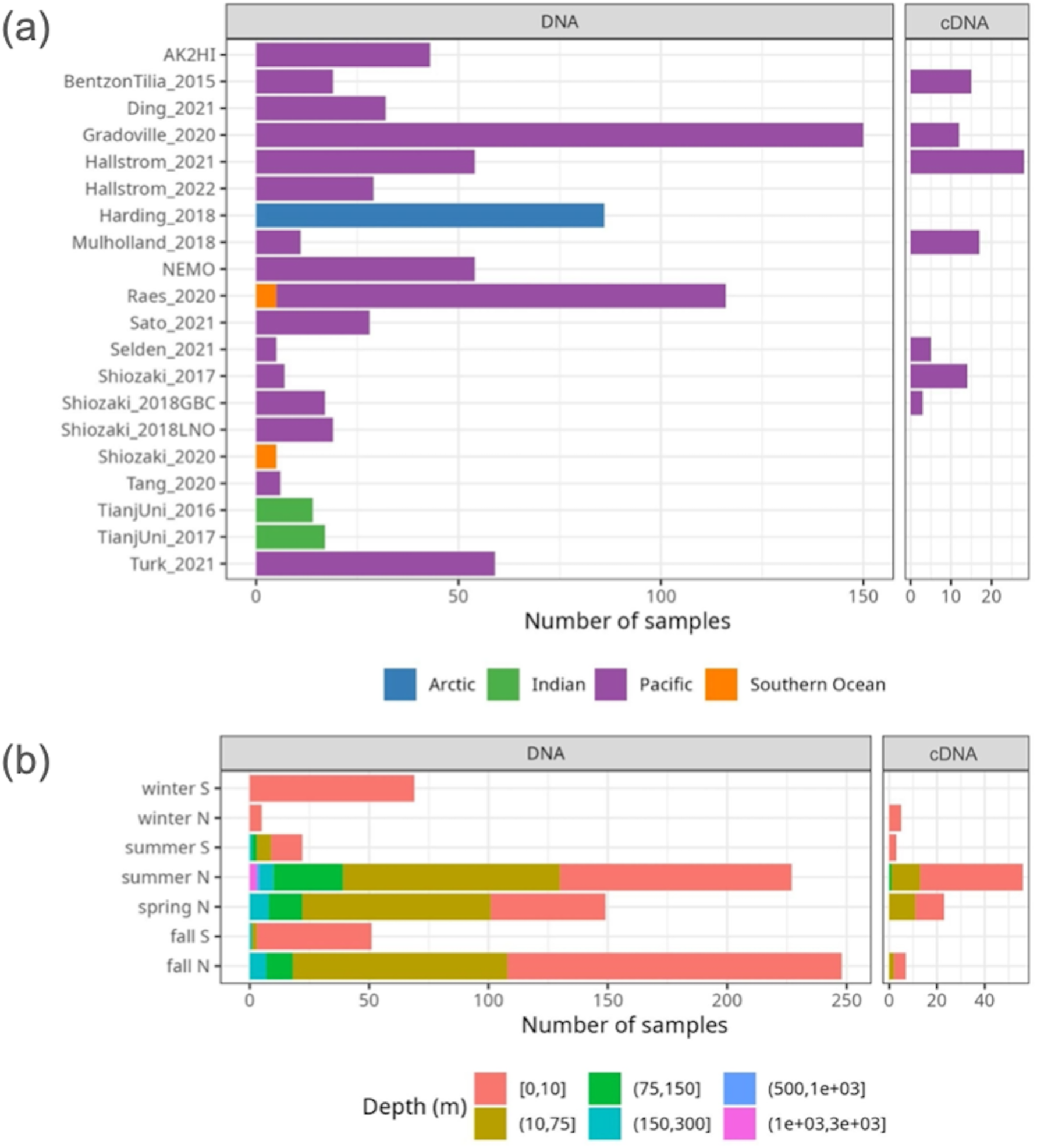
Samples in the *nifH* ASV database by collection location, season, and amplicon type. The number of samples from each study are shown by ocean and study (a), and by the collection season, hemisphere, and depth (b). For both panels the amplicon type (DNA or cDNA) is shown, but *x* axis scales differ between (a) and (b). See Table 1 for citations for the studies in (a). For (b) there were no samples collected between 500—1000 m.

Under-sampled regions include the Eastern South Pacific (n=6) and the Western Indian Ocean (n=0) (Figs. 2, 5a, and 6a). Only two studies originated from the Indian Ocean, a unique environment with intense weather and shifting circulation patterns that include monsoon seasons and upwelling conditions that will require much greater sampling coverage to capture diazotroph biogeography. No South Atlantic samples were found during compilation that met the criteria for inclusion in the *nifH* ASV database, though there are several studies from this region (Supplementary Table 1). Most Atlantic Ocean samples were coastal and from the North Atlantic. Thus, the Atlantic subtropical gyres, which are known to host diverse diazotrophs (Langlois et al., 2005), are underrepresented by *nifH* amplicon data (Fig. 2).

Tropical and subtropical regions, often associated with high temperatures and low nutrients, are highly represented in the database (Figs. 2 and 5a). This likely influenced the ranges of environmental variables with most samples in the database originating from locations with SST above 15 °C and PO_43-_ below 0.5 µmol L^-1^ (Figs. 5c and 5d). Northern hemisphere samples were collected in all seasons, though fewer from the winter. In contrast, most southern hemisphere samples were collected in the winter and fall (Fig. 6b). While most DNA samples are from the euphotic zone (Fig. 6b), cDNA samples are almost exclusively from the euphotic zone, and mainly from the northern hemisphere during the spring and summer (Fig. 6b), indicating an incomplete picture of diazotroph activity.

The disproportionate spatial and seasonal coverage between hemispheres in the *nifH* ASV database mirrors collection biases in other N_2_ fixation metrics including: N_2_ fixation rate measurements; diazotroph cell counts; and *nifH* qPCR data, which are heavily sourced from the North Atlantic (Shao et al., 2023) or, when targeting NCDs, also the North Pacific (Turk-Kubo et al., 2022). These biases underscore the need for future work in understudied regions and seasons.

### 3.3 Study-specific patterns in global diazotroph assemblages in the DNA dataset

To demonstrate how the *nifH* ASV database can be used, a subset of the data was created that comprised of all DNA samples (89.1 % of the total dataset; Fig. 7) and referred to herein as the “DNA dataset”. Samples derived from cDNA (n=94; Fig. 6) were removed. Replicate samples (n=256) or those with multiple size fractions (n=142) were combined by averaging across replicates or size fractions. This reduced the number of DNA samples to 711 and the total number of reads in the count table to 30.0 million from 34.4 million.

**Figure 7.**
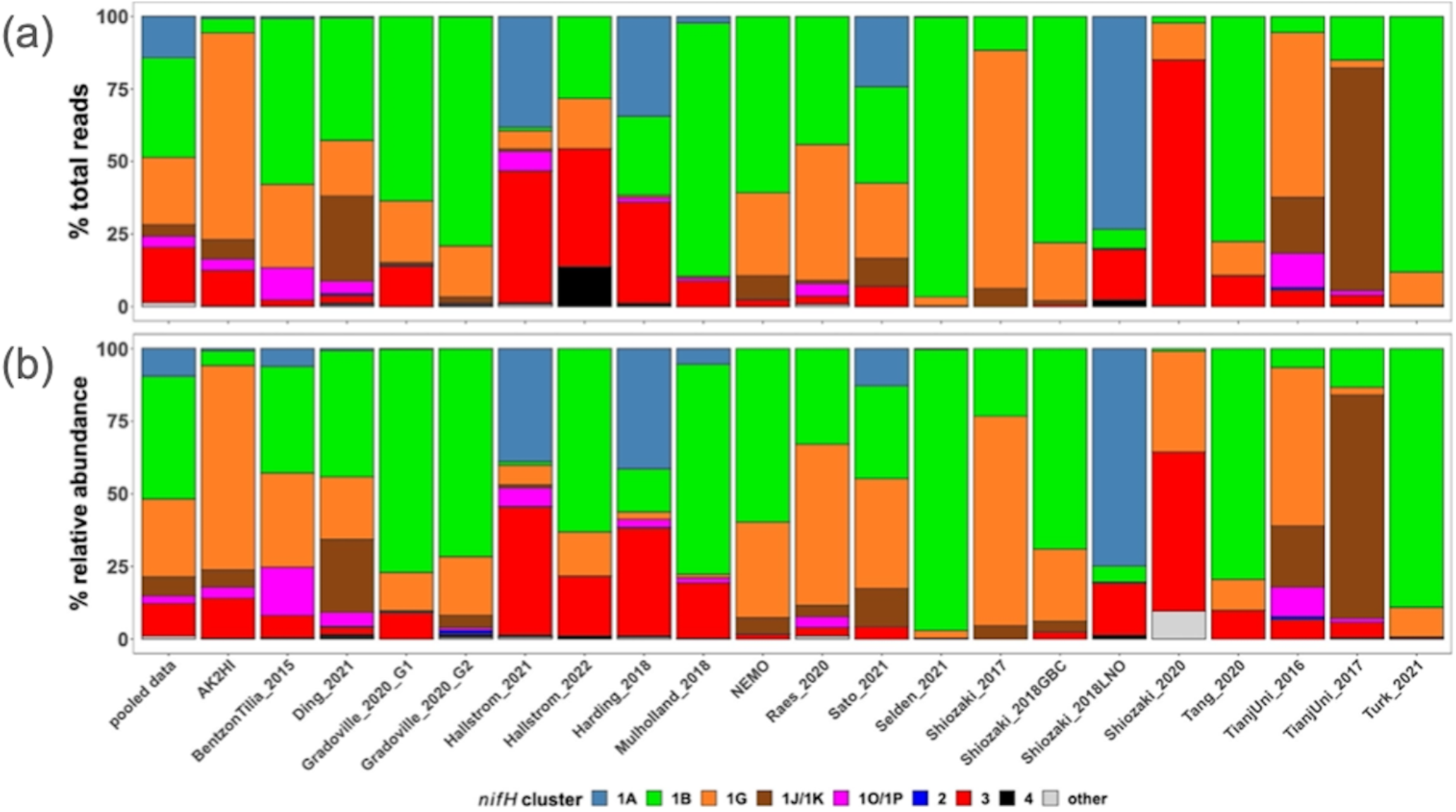
Study-specific diazotroph assemblage patterns in the DNA dataset. The percentage of (a) total reads and (b) relative abundance over the DNA dataset for each major *nifH* cluster. The first column of each panel (‘pooled data’) uses all the compiled data while each subsequent column only uses data from the indicated study. Colors represent different *nifH* subclusters; ‘other’ are the remaining *nifH* clusters.

As demonstrated in a previous global analysis of diazotroph assemblages (Farnelid et al., 2011), cyanobacterial sequences (cluster 1B) dominate the samples, making up 34 % and 42 % of the total reads and relative abundance, respectively (Fig. 7). Though photosynthetic cyanobacteria would be expected to thrive in euphotic waters, NCDs are also widespread in the ocean surface (Langlois et al., 2005; Delmont et al., 2018; Delmont et al., 2022; Pierella Karlusich et al., 2021; Turk-Kubo et al., 2022). Indeed, among the NCDs, γ-proteobacteria (*nifH* cluster 1G) were the most prevalent, comprising ca. 23 % of total reads and 27 % of relative abundance, while δ-proteobacteria (clusters 1A and 3) accounted for 33 % of total reads and 21 % of relative abundance of the DNA dataset (Fig. 7). Less prominent clusters 1J/1K (α- and β -proteobacteria) and 1O/1P (γ-/β- proteobacteria and Deferribacteres) were ca. 4 % and 6 % of the reads and 4 % and 3 % of the relative abundance, respectively. The remaining ASVs comprised <1.5 % of the total reads and relative abundances and came from clusters associated with nitrogenases that do not use iron (e.g. cluster 2) or that are uncharacterized (cluster 4) (Fig. 7).

Cluster 1B (cyanobacteria) were generally high in individual studies across the *nifH* DNA dataset, comprising ≥25 % of the relative abundance community in two-thirds of the studies (Fig. 7), which is the highest of any cluster. Studies carried out in polar regions (Harding_2018, Shiozaki_2018LNO, Shiozaki_2020) and the Indian Ocean (TianjUni_2016 and TianjUni_2017) were distinct from this pattern, with low relative abundances of cluster 1B. Instead, Arctic studies had high relative abundances of cluster 1A and 3 (both primarily comprised of δ-proteobacteria) and while clusters 1J/1K (α- and β-proteobacteria) and 1O/1P (δ-/β-proteobacteria and Deferribacteres) were the predominate groups in the Indian Ocean.

The second most abundant group was the cluster 1G (γ-proteobacteria), making up ca. 25 % of the total reads across the DNA dataset, with study-specific relative abundances greater than 25 % in eight out of 21 studies (Fig. 7). Members of this group were often found at high relative abundances in Pacific Ocean studies (AK2HI, NEMO, Raes_2020, Sato_2021, Shiozaki_2017), as well as in other ocean regions including the Atlantic (BentzonTilla_2015), Indian (TianjUni_2016) and Southern Ocean (Shiozaki_2020). The notable exception is in Arctic studies, where cluster 1G was almost absent (Fig. 7).

In several studies, including BentzonTillia_2015, Hallstrom_2021, Mulholland_2018, Selden_2021, Tang_2020, and Hallstrom_2022, diazotroph assemblages had high relative abundances of putative δ-proteobacteria (clusters 1A and 3), reflecting possibly a coastal/shelf or upwelling signature (Figs. 2 and 7). The only study with samples primarily from the Southern Ocean (Shiozaki_2020) was also the only study with a large portion of *nifH* cluster 1E (*Bacillota*).

#### 3.3.2 Emerging patterns in global diazotroph assemblages across the DNA dataset

The *nifH* ASV database enables new analyses of global diazotroph biogeography in the context of environmental parameters, through co-localization with satellite and model outputs publicly available through CMAP (Ashkezari et al., 2021). To demonstrate the utility of the *nifH* ASV database, we present here patterns in relative abundances of *nifH* clusters across absolute latitude and SST in the DNA dataset. Cosmopolitan distributions were evident for γ-proteobacterial (1G) and cyanobacterial diazotrophs (1B; Fig. 8a), corroborating and extending previous findings (Farnelid et al., 2011; Shao and Luo, 2022; Halm et al., 2012; Fernandez et al., 2011; Löscher et al., 2014; Cheung et al., 2016). At low to mid latitudes, γ- proteobacterial (1G) diazotrophs generally had high relative abundances and were often the dominant taxa when present. However, they declined within the gyre regions, ranging between ∼25–50 % of the population when present, while cyanobacterial diazotrophs (1B) increased and became dominant in the subtropical gyres (Fig. 8a). Notably, cluster 1G diazotrophs reached high relative abundances in each transitional zone, before mainly disappearing at latitudes above 56° (Fig. 8a). However, as mentioned previously, sampling bias likely plays a large role at these higher latitudes where the number of studies and samples are sparse (Figs. 2 and 5).

**Figure 8:**
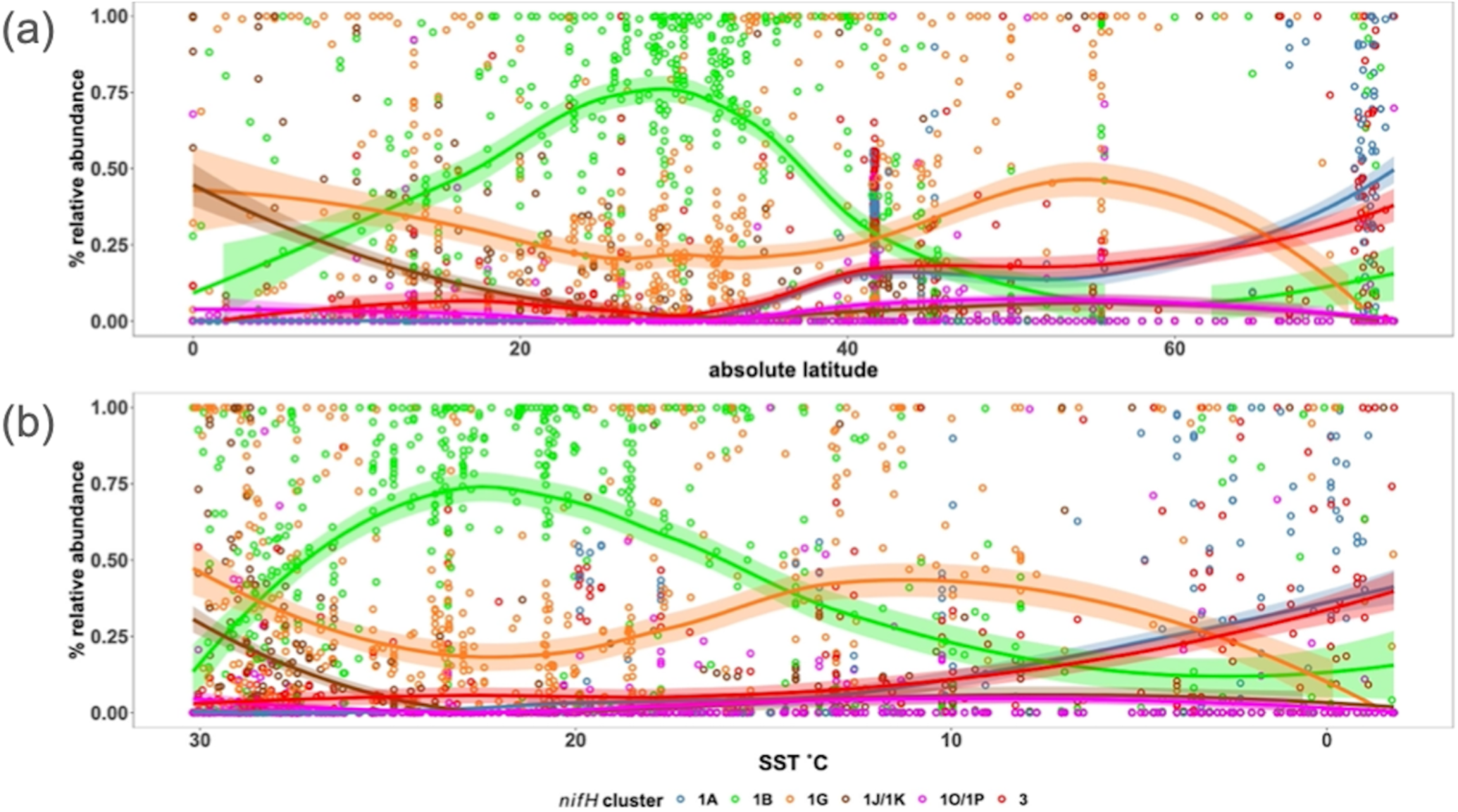
Global distribution of major *nifH* clusters in the DNA dataset. The relative abundance of *nifH* genes for each major *nifH* cluster from every sample compiled in the DNA dataset versus (a) absolute latitudinal and (b) SST. Smoothing averages (lines) were calculated using local polynomial regression fitting (LOESS) with 95% confidence intervals (translucent colored areas). Each color represents a different *nifH* cluster. SST in (b) is from warmest to coolest temperatures to show that trends are similar to those in (a).

Clusters 1B and 1G were both detected over the full range of SST (approximately -2–30 °C) but peaks in their relative abundances occurred in distinct SST ranges. Cyanobacterial diazotrophs had multiple peaks in relative abundance in waters >18 °C underscoring their dominance in tropical gyre regions (Fig. 8b). The 1G cluster also spanned the entire temperature spectrum but had notably higher presence and relative abundance above SSTs of 8 °C and 11 °C, respectively (Fig. 8b). The overlap between 1G and 1B has been reported previously, however the factors controlling this are unknown (Moisander et al., 2014; Shiozaki et al., 2017; Shiozaki et al., 2018b; Liu et al., 2020; Tang et al., 2020; Messer et al., 2015).

δ-proteobacterial diazotrophs (clusters 1A and 3) were generally found in cooler, higher latitude waters. Notably, both clusters 1A and 3 were mainly found below ∼10°C (Fig. 8b). δ-proteobacteria associated with cluster 1A were generally found at latitudes >32° and reached maximum relative abundances near the poles, including in the Beaufort Sea, the highest latitude region surveyed (72°; Figs. 2, 5, and 8a). The vast majority of cluster 1A δ-proteobacteria were found at SST ≤5 °C (Fig. 8b). Though cluster 3 and 1A distributions were similar, cluster 3 showed broader spatial and temperature ranges, with consistent but low relative abundances in the subtropics and tropics (Fig. 8).

In contrast, the relative abundances of cluster 1J/1K and 1O/1P diazotrophs declined as SST decreased and latitude increased, becoming rare at higher latitudes (Fig 8). The highest relative abundances for these clusters were observed near the equator, and in some cases, comprised 100% of the diazotroph assemblage in high SST, tropical samples. These patterns suggest that temperature was an important factor controlling the narrow SST band (≥26 °C) clusters 1J/1K and 1O/1P occupied, establishing them as the *nifH* clusters with the smallest geographic range in the *nifH* ASV database (Fig. 8).

### 3.4 Limits and caveats to interpreting *nifH* amplicon data

The PCR amplification of the *nifH* gene and its transcripts has been vital in advancing the knowledge of diazotroph ecology due to its high sensitivity, detecting diazotrophs at abundances that are often orders of magnitude lower than other marine microbes. This approach has facilitated the discovery of many novel diazotrophs, and provided the first evidence of the widespread distribution of unicellular diazotrophs throughout the open oceans (Falcon et al., 2004; Falcon et al., 2002; Zehr et al., 1998; Zehr et al., 2001). Advances in HTS technologies have revealed diverse diazotrophic assemblages, including the ubiquitously distributed NCDs (Turk-Kubo et al., 2014; Shiozaki et al., 2017; Raes et al., 2020). These discoveries have fostered a new perspective of global diazotrophic ecology (Zehr and Capone, 2020), improved our models of diazotrophic distributions and global N fixation rates (Tang et al., 2019) and will continue to drive new research questions.

However, interpreting *nifH* PCR-based data requires the consideration of several important caveats. Diazotrophs constitute a small fraction of the total microbial community, and thus often require numerous PCR cycles in conjunction with nested PCR for detection. Increasing the number of cycles can exacerbate known amplification biases (Turk et al., 2011) and increase the likelihood of detecting contaminant sequences (Zehr et al., 2003). Strategies to mitigate and assess contamination exist, e.g., by employing ultrafiltration of reagents and including blanks at different stages of the sampling and sequencing process (Bostrom et al., 2007; Farnelid et al., 2011; Blais et al., 2012; Moisander et al., 2014; Langlois et al., 2015; Fernandez-Mendez et al., 2016; Cheung et al., 2021), but such strategies have not been universally adopted. Additionally, relative abundances of PCR amplicons cannot easily be related to absolute abundances. For example, the relative abundance of a taxon can change even if its absolute abundance remains constant, or the relative abundance can remain constant despite changes in the total assemblage size. Moreover, the complexity of the diazotroph assemblage can, if the HTS sequencing depth is insufficient, cause rare ASVs to go undetected, or have relative abundances which are too low to interpret.

Primary objectives in studying marine diazotrophic populations include understanding the contribution of each group to N_2_ fixation, the factors influencing their activity, and their global distributions. The relative abundances of *nifH* genes and transcripts estimated by the workflow can point to potentially significant contributors to N_2_ fixation rates. Yet, the presence of *nifH* genes or transcripts does not always correlate with N_2_ fixation rates (e.g. (Gradoville et al., 2017)). This underscores the need for cell-specific rates to better constrain N_2_ fixation, the assemblages driving given rates, and the taxa-specific regulatory factors of N_2_ fixation to better constrain global biogeochemical modeling.

Various methods are available to target specific diazotroph taxa over space and time (e.g. qPCR/ddPCR, fluorescent in situ hybridization (FISH)-based methods). Universal PCR assays, e.g., those used in the studies compiled here (nifH1-4), are an important complement because they better capture the overall diversity of the diazotrophic assemblage. Unlike primers designed for specific sequences, universal primers can amplify unknown or ambiguous sequences, enabling the discovery of genetic diversity. This includes microdiversity, where sequences show subtle variations from known ones, or even identifying entirely novel taxa. Primers specific to novel sequences can then be developed for use in the mentioned quantitative methods, enabling experiments to characterize the growth, activity, and controlling factors/dynamics of putative diazotrophs growth.

Tools like RT-qPCR, where transcript abundances are assessed directly, or FISH-based methods where single-cells are identified for cell-specific analysis, provide complementary perspectives into the activities of putative diazotrophs. Enumerating diazotrophs using techniques like these can help standardize the relative abundances associated with amplicon sequencing via matching taxa across each method. By assessing diversity and abundance simultaneously, major players can potentially be identified and monitored.

Through genome reconstruction, ’omics studies can enhance the characterization of putative diazotroph amplicon sequences by providing a robust suite of associated genetic data, e.g., taxonomic, phylogenetic, and metabolic. Previous studies have led to the assembly of dozens of diazotrophic genomes (Delmont et al., 2022; Delmont et al., 2018). However, ’omics methods often require massive amounts of data to detect rare community members, and linking genes of interest to other genomic information, e.g., taxonomy, remains quite difficult. Gene-specific models are also required to retrieve diazotrophic information and these models can benefit greatly from the high quality diazotrophic sequences of the *nifH* ASV database. In summary, the complementary perspectives afforded by the methods just described should all be used to obtain robust insights into diazotrophic assemblages.

## 4 Data Availability

The *nifH* ASV database is freely available in Figshare (https://doi.org/10.6084/m9.figshare.23795943.v1; Morando et al., 2024). HTS datasets for the 21 studies in the database can be obtained from the NCBI Sequence Read Archive using the NCBI BioProject accessions in Table 1.

## 5 Code Availability

The workflow used to generate the *nifH* ASV database is freely available in two GitHub repositories, one for the DADA2 *nifH* pipeline (https://github.com/jdmagasin/nifH_amplicons_DADA2) and one for the post-pipeline stages (https://github.com/jdmagasin/nifH-ASV-workflow).

## 6 Conclusion

The workflow and *nifH* ASV database represent a significant step towards a unified framework that facilitates cross-study comparisons of marine diazotroph diversity and biogeography. Furthermore, they could guide future research, including cruise planning, e.g., focusing more on the southern hemisphere and areas outside of the tropics, and molecular assay development, e.g., assays to characterize NCDs for single-cell activity rates.

To demonstrate the utility of our framework, the DNA dataset was used to identify potentially important ASVs and diazotrophic groups, establishing global biogeographic patterns from this aggregated amplicon data. Cyanobacteria were the dominant diazotrophic group, but cumulatively the NCDs made up more than half of the total data. Distinct latitudinal patterns were seen among these major diazotrophic groups, with NCDs (clusters 1G, 1J/K, 1O/1P, 1A, and 3) having a greater contribution to relative abundances near the equator and at higher latitudes, while cyanobacteria (1B) comprised a majority of the diazotroph assemblage in the subtropics. SST appeared to restrict and differentiate the biogeography of clusters 1J/1K and 1O/1P (warm tropics/subtropics) from clusters 3 and 1A (cool, high latitude waters), but did not play as large of a role for the biogeography of clusters 1B and 1G.

We provide the workflow and database for future investigations into the ecological factors driving global diazotrophic biogeography and responses to a changing climate. Ultimately, we hope that insights derived from the use of our framework will inform global biogeochemical models and improve predictions of future assemblages.

## 7 Author Contributions

KTK and MM designed the study with input from SC and MMM. JM created and optimized the DADA2 pipeline for *nifH* amplicon analyses. JM and MM developed the post-pipeline workflow. MM and JM compiled the database, retrieved environmental data from CMAP, and analyzed the database. MM, JM and KTK wrote the manuscript with input from MMM, SC, and JPZ.

## Acknowledgements

We gratefully acknowledge Mohammad Ashkezari and the Simons CMAP team, Stefan Green (Rush University) and the DNA genomics core at University of Illinois at Chicago, Irina Shilova, Julie Robidart and Grace Reed for NEMO sampling and sample processing, and Angelicque White (University of Hawaii, Manoa) and Mary R. Gradoville (Columbia River Inter- Tribal Fish Commission) for AK2HI sampling and sample processing. We would like to thank the authors who directly provided access to sequences: Takuhei Shiozaki (Shiozaki_2017 and Shiozaki_2018GBC) and Jun Sun, Changling Ding, and Chao Wu (TianjUni_2016). This work was supported by grants from the National Science Foundation to KTK (OCE-2023498) and the Simons Foundation to JPZ (Simons Collaboration on Ocean Processes and Ecology, Award ID 724220).

## Supplementary Information

**Supplementary Figure 1. ASV annotations.** The Venn diagram summarizes annotations assigned to 7931 ASVs during the AnnotateAuids stage of the workflow (Fig. 1). Numbers indicate how many ASVs received each type of annotation. Of the 9416 ASVs from the preceding workflow stage, FilterAuids, only the 7931 ASVs shown received annotations.

**Supplementary Table 1. Compiled *nifH* amplicon studies.** Information on all studies compiled to generate the *nifH* ASV database, as well as studies that were not ultimately included and the reasons for this. This comprises the study ID used to refer to each dataset, the number of samples, SRA ID, a reference to each publication and its corresponding DOI.

**Supplementary Table 2. CMAP variables.** A brief description of the environmental variables retrieved from the CMAP data portal.

