## Supplementary figures and images for "Global biogeography of N_2_-fixing microbes: *nifH* amplicon database and analytics workflow"

### figS1.png

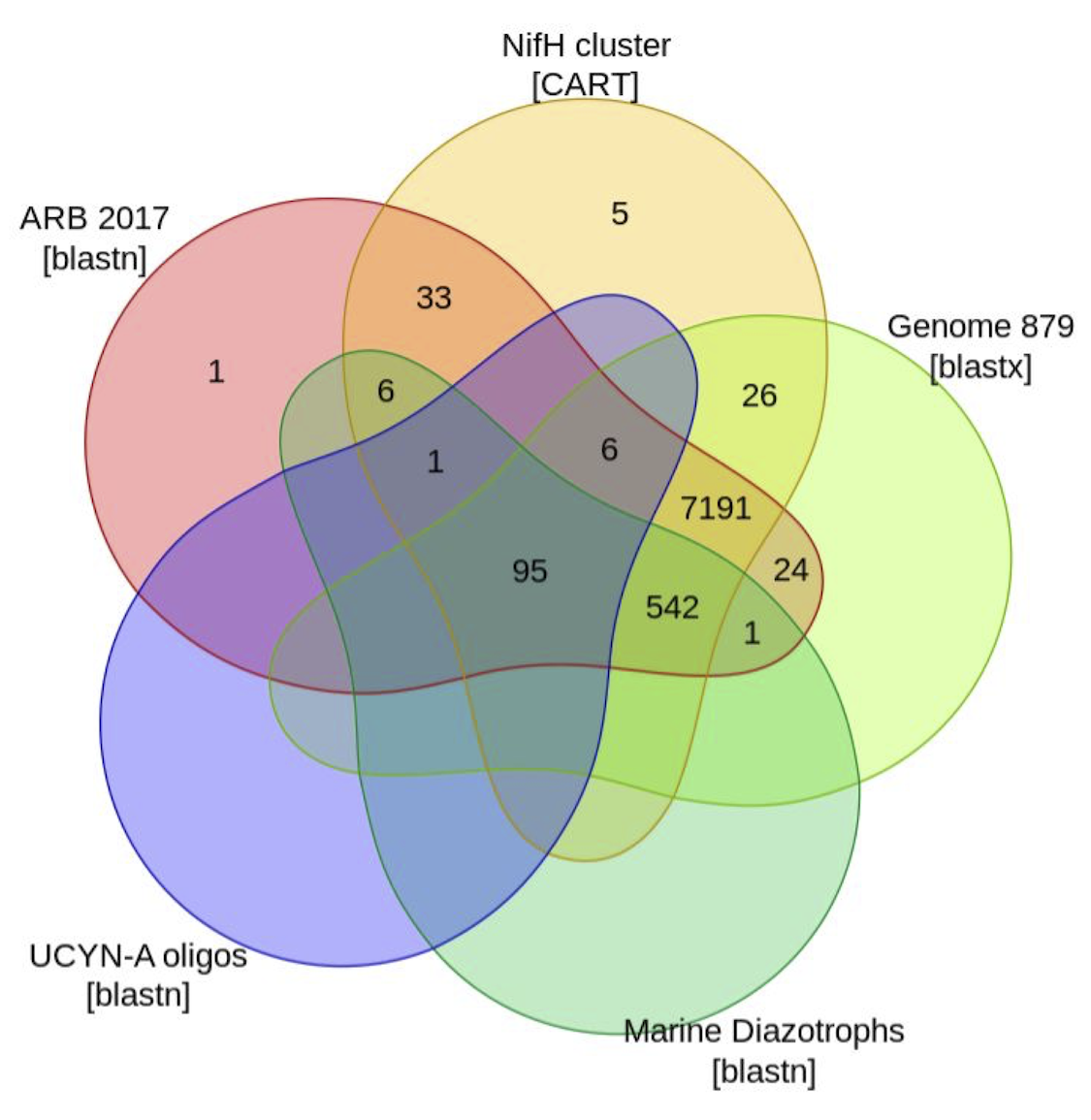
